# Persistent laterality in soaring flight is elicited by demand

**DOI:** 10.1101/2025.09.24.678238

**Authors:** Kamran Safi, Ellen Y. Ye, Andrea Flack, Martin U. Grüebler, Arne Hegemann, Petra Sumasgutner, Patrik Byholm, Elham Nourani

**Author notes:** Corresponding author: Kamran Safi.

## Abstract

Efficient neural processing for executing difficult behavioural tasks is achieved through lateralisation, the favouring of one side. Flight, however, strongly selects for bilateral symmetry, and a persistent turning-side bias could be in theory costly. How soaring birds reconcile lateralised efficiency with symmetric flight is unresolved. We followed 248 individuals of three soaring species, the European honey buzzard, the golden eagle and the white stork, from their first flight for a median of 1.3, 2.2 and 1.7 years per individual respectively, through more than 1.5 million circling manoeuvres. We modelled the turning side of every circle as a difficulty-conditioned process with individual random effects. Laterality was individual: each bird held a stable, repeatable preferred side. Side-bias in circling widened about threefold from the easiest to the hardest circles, and on the days a bird visibly favoured one side, that side was its own seven times in ten. The bias was established early and changed little with age, and its mechanical trace, while present on a single manoeuvre, did not grow into a measurable daily or seasonal cost. Across the three species, lateralisation and symmetric flight coexist: the bias stays faint until difficulty draws it out, yet remains too small to compound.

## Introduction

Behavioural lateralisation—favouring one side when executing a complex task—is a hallmark of efficient neural processing and is considered ubiquitous across the animal kingdom [1, 2]. Much of its advantage rests on the idea that allocating a task to one brain hemisphere, and hence executing it with a side-bias, improves performance [3], streamlining cognition and action from predator detection in chickens to handedness in humans [2].

Even though laterality and hemispheric asymmetry can provide advantages through elevated efficiency at an individual level, they cannot explain why only about one person in ten is left-handed [4]. Individual-level and population-level laterality are distinct: an individual-level bias is a preference a given animal holds consistently, which may point either way and need not be shared with others, whereas a population-level bias is the alignment of those preferences across individuals. Laterality at population level can arise either through ontogenetic and develop-mental pathways that predetermine a direction for asymmetry, or as an emergent property of collectives where assortment of asymmetrical behavioural or morphological characteristics leads to an increased individual fitness benefit [5, 6].

A striking example of developmentally determined population-level lateralisation is the visual asymmetry in birds. Positioned asymmetrically in the egg, bird embryos have their right eye directed towards the translucent shell, and the resulting differential light exposure produces consistent functional asymmetries [2, 7–9], giving rise to a performance difference between the left and right hemispheres and eyes that emerges after hatching. This developmental mechanism thus underlies performance differences in chickens (and other avian species), where non-lateralised individuals make more mistakes than lateralised individuals in spotting predators while feeding, a potentially fatal trade-off between vigilance and the ability to distinguish grit from grain [3]. Lateralisation depends on the task, mainly on its difficulty: easy or well-practised tasks are handled bilaterally, whereas difficult or novel tasks revert to lateralised, hemisphere-specific processing and execution [10, 11]. It is suggested that if both hemispheres of the brain processed information in parallel, the efficiency of the neurological task would be reduced due to the higher cognitive redundancy required [6].

Lateralisation, despite its neurological and motor benefits, creates, however, a dilemma when it comes to difficult tasks that actually require coordinated bilateral execution, because it reduces an animal’s ability to coordinate both sides equally well. Flight is among the most demanding and functionally constrained motor behaviours in vertebrates, and it is where the requirement for symmetry seems most acute: aerodynamic performance is strongly selected for bilateral symmetry, and even slight asymmetry in flight-relevant morphology carries a measurable cost [12]. Morphological symmetry does not, however, preclude a functional side-bias. Empirically, laterality in flight remains little explored, and given centuries of ornithological observation, its near-absence from both the scientific and popular literature is striking. While birds are lateralised in many contexts, from visual processing to footedness [2, 13], a direct test in flight found no side-bias at either the individual or population level [14]. Where a side-bias in free flight has been found, in budgerigars choosing between two equally wide passages, it varied in direction and magnitude from one individual to the next rather than aligning across the population [15]. Only recently have studies begun to quantify laterality in flight, at the individual [16] and population level [17].

Soaring birds extract energy from thermals and do so by circling within narrow, invisible, and ephemeral updrafts, allowing them to gain height, an energy store they then spend against drag as they glide to their next thermal updraft [18–21]. Extracting energy from a rising pocket of air demands precise and continuously adjusted circling [22] (figure 1*a*), and a bird that favours one turning direction must hold that adjustment against its preference on every circle it flies the other way. Such a bias might therefore carry a cost, borne through the roll, pitch and yaw adjustments of circling flight (figure 1*b*). Because performance in soaring has downstream consequences for migration and survival [23–25], even a small per-circle cost could matter if it were to accumulate. The only published account of directional soaring preference we could find reports ambiguous findings: vultures in East Africa circled anticlockwise in three quarters of observations while vultures in the eastern United States showed no preference, and with no way to identify individuals the author attributed the difference, without evidence, to rotation of the air [26].

**Figure 1:**
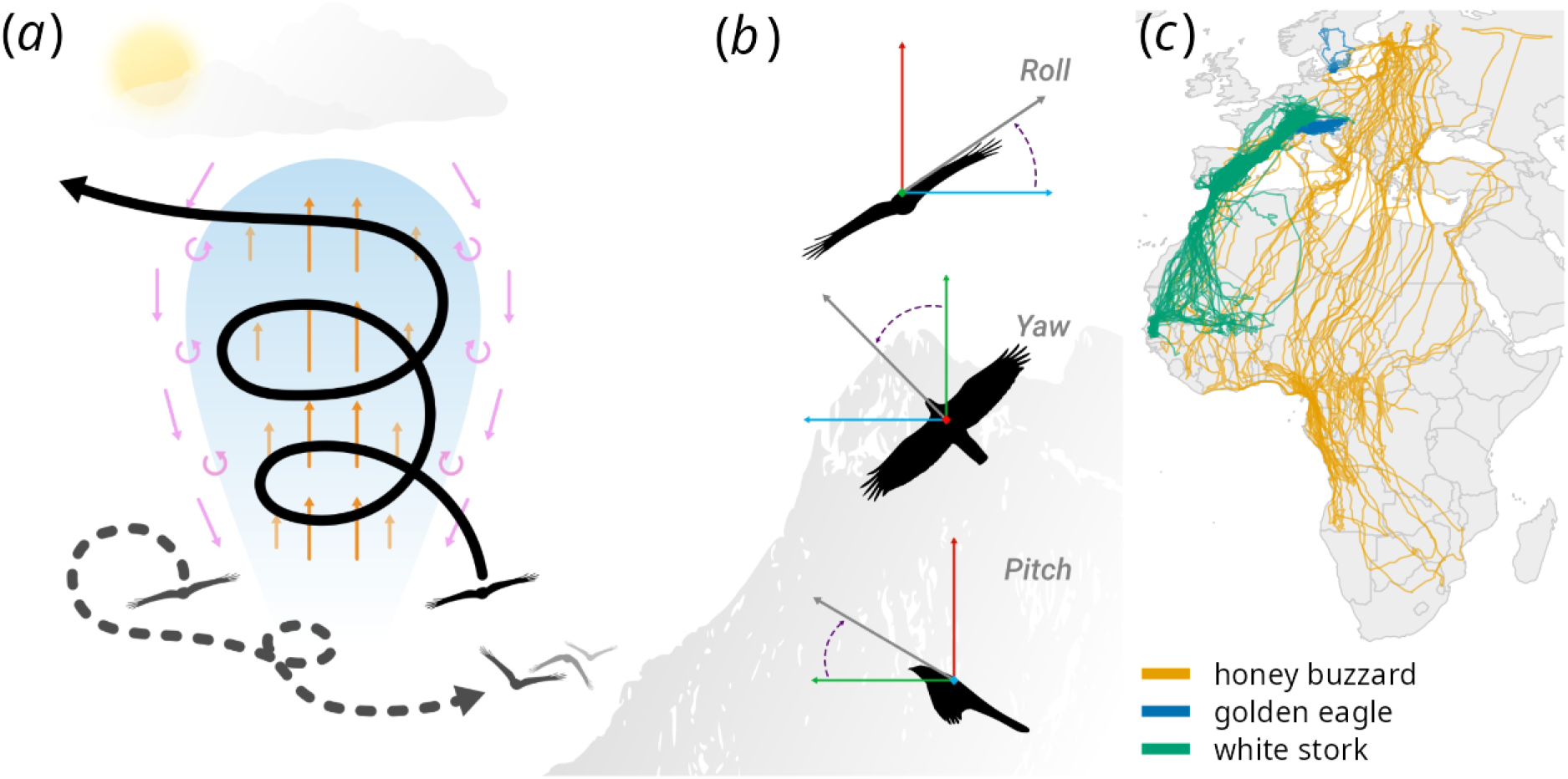
Study systems and thermal soaring. (*a*) Schematic: a soaring bird gains height by circling within a rising thermal updraft (orange arrows) and then glides towards its destination (solid trajectory), while the dashed trajectory shows a bird that does not enter the updraft, and circling requires continuous adjustment of the bird’s orientation. Pink arrows and curls show the downdrafts and turbulent air around the thermal. (*b*) Schematic: the three rotational axes an inertial measurement unit records: roll, yaw and pitch. (*c*) Tracks of the three study species, thinned for display, over Europe and Africa: European honey buzzard (Finland and Switzerland, orange), golden eagle (Sweden and Central European Alps, blue) and white stork (south-western Germany, green).

A population-level bias is either absent from the record or, in the one instance where it was reported, ambiguous. Neither settles the questions as to whether, and if so when, a side-bias appears in soaring flight, how it changes as birds gain experience, and what it potentially costs lateralised individuals. A bias that differs in direction between individuals, or one expressed only when circling is demanding, would leave the population at parity and be invisible to an observer without individual identity. Distinguishing those possibilities requires measuring a preference for each bird, over many thermals, and across a range of demand. We therefore investigate these questions in three soaring species that differ in body size, wing morphology, life history and migratory strategy: the European honey buzzard (*Pernis apivorus*), the golden eagle (*Aquila chrysaetos*) and the white stork (*Ciconia ciconia*) (figure 1*c*). Convergent answers across these disparate systems would point to a general principle of how soaring birds control demanding flight. The answers we obtained are that soaring laterality is an individual, rather than a population-level, trait; that it is elicited by difficulty, being amplified as circling becomes harder; that it is established early and changes little with development or experience thereafter; and that it carries a real but small per-manoeuvre cost that does not measurably accumulate into a daily or seasonal penalty.

## Material and methods

### Study systems and tracking

We analysed individuals of three species carrying solar GPS and IMU (9-axis inertial measurement units: acceleration, gyroscope and magnetometers) tags (e-obs GmbH, Grünwald, Germany), which recorded GPS fixes and, on a subset of birds, high-rate accelerometer and delivered quaternion (rotation angles based on fused acceleration and gyroscope sensors) bursts. We tracked the migratory honey buzzards (HB) from Finland and Switzerland, the resident golden eagles (GE) in the Central European Alps and southern Sweden, and the migratory white storks (WS) from south-western Germany (figure 1*c*). All birds were tagged as nestlings and were tracked from their first flights onward. From 354 tagged birds, 339 remained after a-priori exclusions, which removed monitoring and rehabilitation animals, wrong-population and tag-swap cases. The longitudinal requirement (at least 180 days of GPS, or IMU circles) applies after cleaning and takes the cohort to 196 (electronic supplementary material, hereafter ESM, section S1). Sample size differs among analyses because each uses every bird with sufficient data for it. The cohorts below, and the number of birds behind every reported estimate, are listed in ESM, section S1 and tables S1 and S2. In the analyses of laterality as a difficulty-conditioned response (see below) within circles, namely the per-individual odds, its directional repeatability and its developmental change, we included every individual with sufficient GPS-derived valid circling data: a total of 248 birds (37 HB, 76 GE and 135 WS). Analyses that describe the cohort and summarise laterality at the population level required longitudinal coverage in order to treat a bird as a persistent unit across its annual cycle. For these, we used the 196 birds with valid GPS circling data (37 HB, 75 GE and 84 WS) that also contributed IMU circles or were tracked by GPS for at least 180 days. The strictly IMU-based analyses used 122 IMU-instrumented birds (37 HB, 27 GE and 58 WS). Together the analysed circles exceeded 1.5 million circles flown, and the three study systems are summarised in ESM, table S1.

### Detecting circling and its difficulty

From cleaned GPS tracks [27], we segmented circling behaviour, each characterised by a net 360 degrees of turning. For every circle we determined the turning side (left or right) and a mechanical difficulty axis, the turn rate. The unit is the completed circle, so the analysis begins once a bird is thermalling. We derived the coordinated-turn relation, which is equivalent to the bank angle the bird must hold, so that tighter, harder circles correspond to higher difficulty [28]. We annotated each circle with wind, gust and updraft covariates (ESM, section S8) derived at high spatial and temporal resolution from the birds’ movement recorded by high-resolution GPS, and interpolated to the timestamp of the IMU sensor data where the IMU and GPS were asynchronous (based on [29]). Life stages (post-fledging, migration and wintering) were assigned by an independent latitude-based state machine applied to the GPS track, so that analyses could be conditioned on life stage (ESM, section S6).

### The Inertial Measurement Unit

In an IMU, gyroscopes measure angular velocity, whereas accelerometers measure specific force. During a steady turn, centripetal acceleration tilts the apparent-gravity vector away from true vertical, so an attitude estimator that uses it as a gravity reference cannot recover the true bank angle [30]. In our data the IMU-derived absolute bank after correction for sensor mounting orientation was about 6 to 7 degrees against the 23 degrees the circling kinematics require (ESM, section S2), so we do not use absolute IMU bank in any model. The direction of laterality is taken from the GPS turning sense, validated against the IMU yaw, with which it agrees on 98.4 to 99.8 per cent of concurrent circles (ESM, section S3). Bank, load factor and sink rate are taken kinematically from the track, and the IMU contributes only in-flight instability, the standard deviation of roll within each second, and vectorial dynamic body acceleration (VeDBA), a proxy for the dynamic component of effort [31, 32]. One tag with a demonstrably faulty orientation sensor was excluded from the IMU analyses.

### The modelled entities

Quantifying individual laterality from a limited number of observations per bird is statistically demanding. Summarising each individual by a raw laterality index and then classifying it against a threshold is commonplace in the field, but discards uncertainty and treats individuals with very different sample sizes as equivalent, even though their index estimates do not carry the same variance [33]. We therefore model the binary turning side of every circling bout directly, as a difficulty-conditioned binomial process with individual random effects, and we express laterality as a per-individual odds (on a log-scale) rather than as a pooled or thresholded index. This choice recovers the quantity the biological question requires, and it removes several artefacts that follow from a pooled approach, for example based on a laterality index, which is by construction highly susceptible to small-sample bias.

For every question, the unit of analysis is the individual, and the response is the binary turning side of a circle, which we model directly rather than through a pre-computed or thresholded laterality index. A bird’s preferred side is the side it turned to on more than half of its circles, easy and hard together. Because the count ignores difficulty, the slope on difficulty is not an artefact of it. The intercept is, so we report it as a floor and not as a magnitude. The floor analysis, which needs a side defined without the easy circles, takes it from hard circles alone. Because we model the probability that a circle is flown to a bird’s preferred side, laterality here is a graded preference along a continuum rather than a categorical handedness, and a bird classified as lateralised still uses both directions, as observed in many phenomena of lateralisation [34, 35]. We fitted binomial generalised linear mixed models (glmmTMB) with nested individual and thermal ID random effects [36]. Difficulty enters through a floor-plus-gain hinge, defined as the positive part of difficulty above a species-level onset: an intercept captures each bird’s baseline shift from that species-level floor, and a random gain slope captures how strongly the bird amplifies that shift once difficulty exceeds the onset (ESM, section S5). The onset is 27 degrees per second in HB, 15 in GE and 21 in WS. Circles below it are easy, above it hard. One standard deviation of turn rate is 12.7, 9.9 and 8.7 degrees per second. The gain odds ratio, estimated for the population and as a random slope for every individual, is the central modelled parameter of the study. Overdispersion, per-individual shrinkage, spatial (Moran’s I [37]) and within-bird temporal (lag-1) residual independence, AR1 sensitivity, and Benjamini-Hochberg control of multiplicity [38] are treated in ESM, sections S4, S7 and S10, with the two multiplicity ledgers in ESM, tables S3 and S3b. Because inference is over individuals, we report population-level p-values only as a check of parity (ESM, section S7). We fitted all models in a frequentist framework, and give full model specifications, convergence and diagnostics in the model cards of ESM, section S4.

### Costs

We quantified the mechanical signature of the side bias per manoeuvre, namely the in-flight instability asymmetry between the preferred and non-preferred side and the climb and energetic penalty of reversing turn direction. To ask whether this cost accumulates, we tested whether a bird performed worse on the days it circled more often against its preferred side. For every bird and every day we counted the circles flown on the non-preferred side, weighting each circle by its difficulty, and did the same for the preferred side. We then compared each bird with itself: was its performance on days with many wrong-side circles lower than on its other days, holding the amount of preferred-side circling and the total number of circles that day constant? Hard days are hard for every bird, so a poorer performance on wrong-side-heavy days could simply reflect difficult conditions. To separate the two we ran the same comparison in the symmetric birds, those with no detectable side preference, for which the label wrong side is arbitrary and which therefore serve as a placebo, and took the difference between the two: the daily side cost is the gap in the lateralised birds minus the gap in the symmetric birds, a difference-in-differences [39]. We measured daily performance in five ways: the rate at which a bird gained height in thermals (climb rate) and the speed at which it progressed across the landscape (cross-country speed), which we had chosen in advance as the two decisive measures, and three further measures that we report as secondary, the rate at which it lost height while gliding, the straightness of its route and the number of thermals it used per kilometre. Standard errors were clustered by bird (ESM, section S8). We additionally accounted the estimated per-circle deficits deterministically onto each bird’s realised circling durations, to bound the daily time cost (ESM, section S8). All performance results are reported as associations, with difficulty, wind and stage (post-fledging, migration and wintering, or resident) controlled.

## Results

### Laterality is an individual trait, not a population handedness

At the population level, all three species circled close to parity, with only small deviations from 0.5, significant in one of the three (figure 2). White storks (0.496) and honey buzzards (0.514) were indistinguishable from parity, and the golden eagle carried a small but significant rightward bias at the population level (0.530), detectable only at this very large sample size and considered biologically negligible in magnitude. Below the species onset, on easy circles, the population mean was flat in all three species, at parity in the honey buzzard (0.514) and the white stork (0.496) and at the rightward offset in the golden eagle (0.530). Above the onset the three differed: in the honey buzzard the population mean drifted rightward as circles tightened, to 0.58 two standard deviations above the onset (slope 0.14 per SD on the log-odds scale, p = 0.038), because more buzzards lean right than left. In the golden eagle it held its offset and in the white stork it held parity, while the individual curves fanned out in all three species (ESM, sections S4 and S7). In each species, the deviation from 0.5 is small and dwarfed by the between-individual variance, so the population mean is a check of parity, not evidence for population-level handedness.

**Figure 2:**
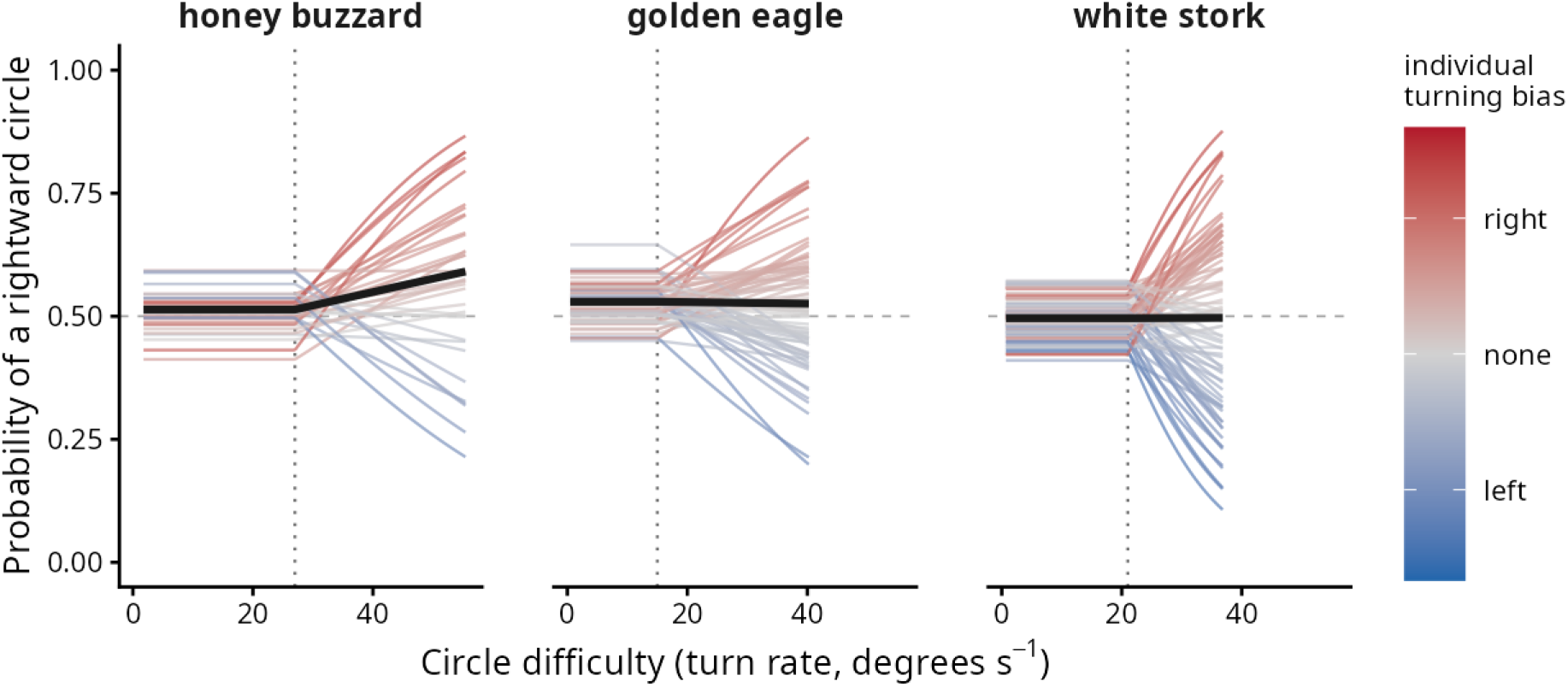
Individual laterality is elicited by difficulty. Fitted per-individual curves from the floor-plus-gain model (Methods), giving the probability of a rightward circle as a function of circle difficulty (turn rate), for each species. Each thin line is one bird, coloured by its own turning bias (blue, left; red, right). Below the species onset (dotted line) all birds sit near parity, the floor. Above it each bird’s preference lifts to its own side, the gain, and the individual curves fan apart. The solid black line is the fitted population mean: flat below the species onset in all three species, and above it rising in the honey buzzard, to 0.58 two standard deviations above the onset because more buzzards lean right than left (population slope 0.14 per SD, p = 0.038), constant at a small but significant rightward offset in the golden eagle, and at parity in the white stork. The panels share one difficulty axis and each species’ curves are drawn over its own observed range. The population probability of a rightward circle is 0.514, 0.530 and 0.496 in the honey buzzard, golden eagle and white stork, the between-individual standard deviation of the baseline preference 0.20, 0.18 and 0.21 on the log-odds scale, and the population gain odds ratio 1.23, 1.11 and 1.25. The gain odds ratio is from the preferred-side model, which uses all 248 birds with valid circles rather than the 196 drawn here.

Individuals, however, were far from symmetric, and the quantity that captures an individual side-bias is the between-individual variance in turning side, not any single bird’s index. That variance is well away from zero in every species: the between-individual standard deviation of the baseline side preference is 0.20 in HB, 0.18 in GE and 0.21 in WS on the log-odds scale (95% confidence interval [CI]: 0.15 to 0.27 HB, 0.15 to 0.23 GE and 0.18 to 0.26 WS), so individuals differ in their preferred side by more than sampling alone would produce. A latent per-circle repeatability of the side was, by contrast, near zero (0.013 HB, 0.013 GE and 0.026 WS), likely because a binary outcome carries a large irreducible latent residual, and because the bias is elicited by difficulty, a repeatability pooled across easy and hard circles is diluted by the many easy events. In hard circles, where the bias is expressed, the individual preference was both substantial and repeatable. A bird’s turning side in the first half of its entire tracking record predicted its side in the second (test-retest correlation 0.62 HB, 0.65 GE and 0.59 WS), and their committed side did not flip from one period of the record to the next more than by chance (serial-persistence p = 0.002 HB, 0.019 GE and 0.001 WS). Each bird therefore carried a repeatable preferred turning side that was expressed when the difficulty of the manoeuvre demands lateral execution, while the population as a whole remains balanced (figure 2; ESM, section S4). This bias is a graded preference, not an exclusive use of one side. On hard circles a bird turned to its preferred side on 0.55 to 0.56 of occasions, on the tightest tenth of its circles on 0.57 to 0.60, and between one bird in five and one in nine (19, 11 and 17 per cent) reached at least a 70:30 preference on those tightest circles. On the days a bird visibly favoured one side when circling was hard, that side was its own on 0.70, 0.67 and 0.72 of days (ESM, section S4). Both directions therefore remained fully available, and individuals differed in how strongly, not in whether, they favoured one side over the other, with about a quarter of birds showing no detectable preference at all (below).

### The bias is elicited by difficulty

Laterality turned out not to be a fixed habit but a conditional response to the difficulty of the manoeuvre. Modelling the turning side as a floor-plus-gain function of circle difficulty, individuals expressed only a faint shift from equal proportion on easy, wide circles (the floor), and amplified it as circles tighten beyond the species onset (the gain). The population gain odds ratio, the amplification of the preferred side per unit of difficulty above the onset, is 1.23 in HB, 1.11 in GE and 1.25 in WS (37, 76 and 135 individuals respectively, 248 in total; all p < 0.001). Estimated bird by bird, the gain odds ratios had medians of 1.15, 1.07 and 1.23 (interquartile ranges 1.00 to 1.51, 0.97 to 1.23 and 0.96 to 1.51), with 73, 66 and 70 per cent of individuals above one. Because each odds ratio is expressed per within-species standard deviation of turn rate, and a given turn rate is a different aerodynamic demand in a fast eagle than in a slower stork, the three magnitudes are not on a common scale. What is comparable is the direction: in all three species the odds of flying to the preferred side rise with difficulty. The three species nonetheless circle at a closely similar median bank of about 23 degrees, so they sample a comparable flight regime (see ESM, section S2). The between-individual spread in turning tendency, measured as the ratio of between-individual standard deviations on the model’s logit scale, fans open roughly threefold from easy to hard circles, as laterality is elicited by difficulty (figure 2). The effect is robust to spatial and temporal autocorrelation, being estimated between birds rather than over autocorrelated circles (ESM, section S7), and it reproduces on the IMU turning signal, which confirms the sign and significance of the difficulty-gain rather than its magnitude, the IMU difficulty axis being shaped by burst length and so not on a common scale with the GPS estimate (ESM, section S4).

The floor is itself a genuine, faint individual lean rather than an artefact: Each bird’s easy-circle lean aligns with its own hard-circle side.

### The bias changes little with development

Laterality in circling proved to be established early in life and changed little thereafter (ESM, section S5). The difficulty-gain steepened significantly with age in honey buzzards (p = 0.013) and white storks (p < 0.001), but not in golden eagles, but in every case the magnitude was negligible: over the entire first year of life the fitted probability of turning to the preferred side on a hard circle rose by only about one percentage point in the buzzard and half a point in the stork, and not at all in the eagle (ESM, figure S6). The side itself was fixed from the earliest recorded flights, and every effect of an experience proxy (number of circles flown) cleaned of confounded effort (age) is indistinguishable from zero in all three species, which also addresses the concern that days since tagging is a coarse experience proxy. Thus, neither the direction of the bias nor its strength changed over the first year.

### The bias carries small per-manoeuvre costs that do not accumulate

The asymmetry left a real but small mechanical trace. In-flight instability, the standard deviation of roll within each second of a circling act (about 2.3, 3.0 and 2.1 degrees per bird in HB, GE and WS), was higher when a bird circled against its preferred side, by 2.2, 2.9 and 0.3 per cent at matched turn rate. The raw side difference, non-preferred minus preferred, was 0.045, 0.059 and −0.002 degrees of roll standard deviation, resolved as the within-bird mean over tens of thousands of paired acts rather than by any single reading (ESM, section S2), significant on the per-bird test in honey buzzards and golden eagles (p = 0.006 and 0.002) but not in white storks (p = 0.23; figure 3). We report this phenomenologically, as greater in-flight instability rather than a demonstrated control deficit. It survives conditioning on within-circle banking variability in the two raptors (ESM, section S2). On the non-preferred side the birds flew at a slightly lower, less settled bank rather than an over-banked one, which we interpret as a reduced-motor-fluency signature (ESM, section S8). This trace does not grow uniformly with difficulty across species, strengthening in the eagle and stork but weakening in the buzzard, so we do not treat growth with difficulty as a mechanistic hallmark, especially given the faint magnitude we recorded in the large data volume analysed.

**Figure 3:**
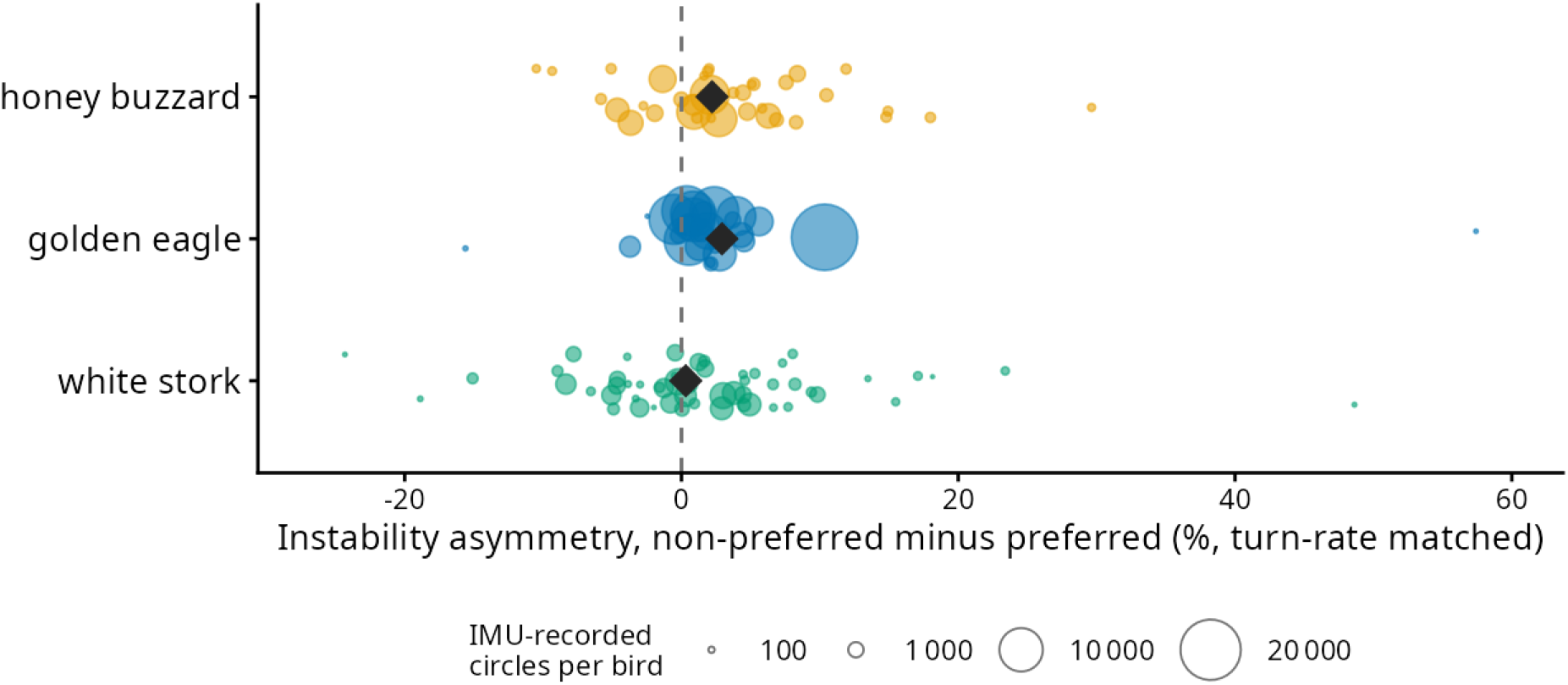
Laterality leaves a real per-manoeuvre instability signature. Each point is one bird’s turn-rate-matched in-flight instability asymmetry, that is the percentage by which its in-flight instability is greater on the non-preferred than on the preferred side. Each point is sized by the number of IMU-recorded circles the bird contributes, so that well-sampled birds carry visual weight and noisy low-sample birds do not. The vertical spread within a row is random jitter that separates overlapping birds and carries no information. The black diamond is the turn-rate-conditioned effect estimated by the per-act model, not the mean of the plotted points. In the two raptors most birds lie right of zero (27 of 37 honey buzzards, 22 of 27 golden eagles), so the non-preferred side is less stable broadly across individuals rather than through a few outliers. In the white stork the split is near even (30 of 54, the paired test including the 54 of 58 IMU storks with IMU-recorded circles on both turning sides; ESM, section S8) and the effect is not significant. The turn-rate-conditioned effect is 2.2, 2.9 and 0.3 per cent, significant on the per-bird test in buzzards and eagles but not in storks (black diamond). This per-manoeuvre cost does not accumulate into a daily or seasonal penalty, which is shown by the null daily difference-in-differences and the deterministic cost accounting in ESM, section S8.

Circling on the non-preferred side carried a climb cost in the eagle alone (per bird, non-preferred minus preferred circles within the same thermal: 0.04 lower in GE, 95% CI 0.01 to 0.05 m s*^−^*^1^). In the two migrants the same contrast did not differ from zero (0.02 lower in HB, from 0.03 lower to 0.00 higher; 0.01 lower in WS, from 0.02 lower to 0.01 higher). Reversing turn direction within a thermal carried a brief penalty: a transient per-event climb-rate deficit (per bird, the circles around a reversal minus steady circles: about 0.12 in HB, 0.29 in GE and 0.16 in WS m s*^−^*^1^) and a within-burst rise in dynamic body acceleration (6.6 in HB, 6.4 in GE and 4.0 in WS per cent, surviving control for gust in the two raptors). The climb deficit is however an association and also a consequence of the external conditions because poor lift both provokes reversals and depresses climb (ESM, section S8).

Not every bird was lateralised: of the individuals with at least 100 hard circles, 8 of 37 honey buzzards, 18 of 71 golden eagles and 33 of 118 white storks circled at parity even on hard circles (ESM, section S4). These symmetric individuals were not less stable fliers than lateralised ones: their in-flight instability at matched turn rate, gust and updraft differed from that of lateralised birds by −0.3, −3.2 and −5.0 per cent (p = 0.73, 1 and 0.33 on the between-bird test; 16, 5 and 7 symmetric birds with IMU data; ESM, section S8). The wrong-side instability was therefore a difference between a bird’s two sides, not a difference in overall stability between lateralised and symmetric birds. The dynamic body acceleration captures the effort of this extra instability and flapping but not the isometric cost of holding a bank (ESM, section S8).

The measured per-manoeuvre cost, however, did not accumulate. Integrated over each bird’s realised flight, the estimated per-circle deficits bound the extra climbing a lateralised bird incurred at about 0.6 per cent of daily climbing time in honey buzzards, 2.0 per cent in golden eagles and 0.4 per cent in white storks. Thus, the deterministic cost of the side bias alone amounts to only a few minutes across an entire autumn migration in the two migratory species (again under the caveat that adverse atmospheric days lead to both more corrections and less distance flyable, ESM, section S8). The daily test found no side cost: on days when a bird circled more often against its preferred side it neither climbed worse nor travelled more slowly, the two measures we had chosen in advance as decisive, in any of the three species, and the confidence intervals bound any daily side penalty to within about 10 per cent of a typical day’s climb rate in the eagle and stork, the buzzard interval being wider (ESM, section S8). Among the secondary daily outcomes, a single nominal effect appeared, a higher glide sink rate on wrong-side-heavy days in honey buzzards (0.18 m s*^−^*^1^ per unit of difficulty-weighted non-preferred circling, p = 0.038), one of five daily measures in the species whose daily test rests on about 37 birds, and it does not survive correction for multiple testing (ESM, section S8 and table S3b). The apparent daily cost seen in raw data was explained entirely by difficulty exposure, in that hard days are simply harder for lateralised and symmetric birds alike. Consistent with this, we could not detect any relationship between laterality and the speed or completion of the first autumn migration in the migratory species (ESM, section S8).

The fraction of hard circles on which a bird expressed its preference differed across life stages, peaking in migration in buzzards and in wintering in storks, with no stage in which it was suppressed (ESM, section S6). Here too, because difficulty is set largely by the environment, our interpretation is that laterality was elicited by conditions rather than chosen.

## Discussion

Across three morphologically, ecologically, and phylogenetically distinct soaring species, the clearest and most general result was that a turning-side bias was elicited by circling difficulty: from a faint individual baseline, birds amplified their preferred side as circling tightened (population gain odds ratio 1.11 to 1.25). This demand-elicitation converged across honey buzzards, golden eagles and white storks and it points to a general feature of how soaring birds meet the mechanical demand of a tight and hard circle. The other properties of the trait were individual rather than population-level: the bias was an individual trait, not a population handedness; it was established early in life and changed little with development in any species; and its mechanical cost, though real, was small and did not accumulate.

The cost of laterality was measurable at the level of individual manoeuvres. On the non-preferred side, flight was measurably less stable in the honey buzzard and the golden eagle (higher in-flight instability by 2.2 and 2.9 per cent), reversing turn direction within a thermal lost climb and raised effort transiently, and golden eagles turning on their wrong side climbed marginally worse. These small, per-manoeuvre costs were not always detected. Specifically, instability asymmetry in storks and the wrong-side climb deficit in the two migrants (honey buzzard and white stork) did not differ from zero.

We did not find any substantial support that these micro-costs accumulated: integrated over each bird’s flight, the side bias alone bounds the extra climbing at roughly two per cent of daily climbing time or less, and the daily difference-in-differences of climb rate and cross-country speed was null in all three species despite the very large number of events analysed here. We could not detect a relationship between laterality and migration speed or completion (based on the 15 HB and 90 WS with a completed first autumn migration and a laterality estimate, see ESM, section S8). However, our tracking under-represented harsh flight conditions such as marginal thermals, strong shear and competition where a cost might be more consequential. Consistent with a low cost, birds continued to turn against their preference on a substantial share of hard circles, an equilibrium a large net penalty would be expected to erode either during ontogeny, with gaining experience, or through natural selection [5, 6, 13, 40].

Not every bird carried a side. About one in four in each species (8 of 37 honey buzzards, 18 of 71 golden eagles, 33 of 118 white storks) circled at parity even where the others expressed their preference. These symmetric birds were no more and no less stable in flight than lateralised ones, and did not do better on a given day for having no preferred side to leave. That neither the lateralised birds within themselves nor lateralised and symmetric birds differed in performance leaves the mechanism open: whatever sets and maintains a side, our data cannot tell whether it is kept because it pays or merely costs very little if anything.

In lateralised behaviours, the involvement of two hemispheres in the execution of a difficult task is proposed to shift with its demand [41, 42] such that two main outcomes over an individual’s lifetime are imaginable. In one, manifestation, the preferential use of one side elicits a life-long learning feedback loop, giving rise to a unilateral improvement that manifests asymmetrical execution through motor and perceptual learning and reinforcement [3, 6]. The more one side is used, the more experienced and efficient the lateral execution becomes [2, 43]. The second, simplification, with an opposite outcome, is that with gaining experience and improvement in executing complex behaviours, the perceived difficulty and complexity decreases, reducing the need for laterality and thus allowing bilateral execution [44, 45]. Laterality in all three species studied here was, however, neither overcome nor diminished with experience. If anything, laterality solidified with age when flight was hard, by about one percentage point over the first year in the buzzard and half a point in the stork, suggesting a manifestation of lateral execution with age. However, our sample cannot establish the origin or the mechanisms maintaining a side-bias. The underlying side-bias was fixed from the earliest recorded flights in our data, and the difficulty-gain steepened with age only marginally in the two migrants, by around one percentage point of preferred-side probability over the first year, while not changing in the eagles. Thus, laterality in soaring seems to be an individually stable early-established trait, not a developing skill, in contrast to the soaring performance it is embedded in, which does improve individually with age and developmental stage [22, 46, 47]. What determines the side preference is not known. Heritability of a lateral preference is reported in a topminnow and in chimpanzees and absent in mice [48]. In humans the genetic models do not account for handedness on their own, and perinatal asymmetries of perception and later social modulation are the candidate environmental influences, with the evidence for either incomplete [49]. In the absence of a genetic pedigree and of detailed records of the environmental conditions during development, our data can only add one constraint: whatever sets the side has acted before the first recorded flight.

Only the golden eagle carried a constant population bias, a rightward preference of about 0.53 at every difficulty. In the honey buzzard the population mean was at parity below the onset and drifted rightward above it, to 0.58 on the tightest circles, a composition effect of a cohort of 37 in which more birds lean right than left. Its detectability is a property of scale: across hundreds of thousands of circles an offset far too small to have been noticed in a century of observation of bird behaviour becomes statistically significant in our extensive data. The bias, while real, is probably biologically negligible, and its emergence from noise reflects the reach of biologging and the ability to follow individual animals over long time periods in high detail, more than a population-level handedness.

We are careful about what these results establish. They concern behavioural side-biases and their ontogenetic and performance correlates. We cannot make any claims about neural lateralisation: a behavioural preference of this kind is as compatible with a peripheral or structural asymmetry as with a cerebral one [50]. Although we find little evidence for meaningful costs and lack any quantitative measure of benefits, the turning preference may still be under selection over the longer term, through fitness and survival, on the balance a species strikes between motor skill and the efficiency of neural processing. The side-bias we document is, moreover, a graded turning preference rather than the near-exclusive limb dominance the term handedness suggests. That is what the measurements show elsewhere too: the degree of handedness is continuous even in humans [50, 51], and categorical classifications of lateral preference have no agreed cut [34, 50]. We reported the performance relationships as associations, with difficulty, wind and life stage controlled. The common-cause confound, in which hard conditions produce both more biased circling and poorer travel, is what the difference-in-differences design is built to separate.

### Pitfalls of thresholded laterality indices

An additional contribution of this study is methodological. Our estimates of individual laterality rest on a continuous model of the turning side rather than on a thresholded index such as the laterality index [52]. Index-based laterality carries a small-sample artefact by design that bears weight beyond our study. An index-based analysis would strongly bias the estimates of the side-biased circling behaviour revealed in this study at the per-individual sample sizes available, despite the exceptionally large overall volume of data. For a given underlying turning probability, indices calculated from few observed turns are substantially more variable than those calculated from many turns. Because the absolute laterality index quantifies the magnitude of departure from an even left–right split, thresholding converts large chance deviations into categorical classifications of lateralisation. Sparse observations may therefore spuriously inflate estimated prevalence and strength of lateralisation [33]. This is especially problematic when the behaviour of interest is rare, difficult to observe, or unequally sampled among individuals, life stages, conditions, or locations: variation in observation effort can then be confounded with apparent individual variation in lateralisation.

We therefore modelled the turning side of each circling event as a binary response in a difficulty-conditioned mixed-effects binomial model with random effects for individual birds. This event-level approach retains the full observation-level structure, rather than reducing each bird’s observations to a thresholded laterality index. Modelling individual binary choices directly retains the sampling process and allows covariates, repeated observations, and dependence structures to be represented explicitly, thereby reducing the risk of conflating sampling variation with evidence for or against individual lateralisation. We offer this approach as a template for a field in which laterality has often been characterised through indices and cut-offs [11, 34, 35, 53, 54], but in which modelling turning side directly can yield individual-level inference with uncertainty propagated through to the final estimates.

### Laterality as a cryptic trait with condition-dependent expression

If a side-bias is latent on easy tasks and elicited only by difficulty, then laterality measured under standard or undemanding conditions may routinely be underestimated or missed, and the more informative question is not whether an animal is lateralised but at what level of demand its bias emerges. Read this way, what we observe is not a bias that is acquired or lost through development but a stable, early-established individual bias whose expression is governed by immediate demand rather than by age: the side is set early and changes little, while its expression is drawn out by difficulty. We speculate that a demand-elicited laterality of this kind may be common in complex motor skills, and that probing it across graded difficulty, rather than at a single operating point, would reveal lateralisation where fixed-condition assays find none. Biologging makes this tractable in free-ranging animals at individual resolution, along a measured gradient of demand, over years of the same individual’s life and with the performance of every manoeuvre recorded alongside it, at sample sizes that expose effects that a century of observation could not, and extends the difficulty-modulation hypothesis from birds in experimental flight tasks [10] to free flight in three unrelated species.

Framed as a conditional and idiosyncratic individual trait that costs little, laterality also becomes a candidate axis of the standing behavioural variation increasingly recognised among individuals [55], and, because it is expressed in the act of moving, of the individual variation in movement itself [56]. A bias that surfaces only under load and does not compound could persist, and differ among individuals, without ever appearing as a population signal or paying an obvious fitness price, so that cryptic individual asymmetries may be far more widespread than the sparse record of overt flight laterality implies. The benefit of being lateralised, where it has been shown, is a benefit under competing demands: weakly lateralised chicks discriminate grain from pebbles as well as lateralised ones and fall behind only when they must also watch for a predator [3, 6]. We measured the motor act alone. Whether symmetric birds fall behind when circling competes with another demand, another bird in the thermal, a predator, a decision about where to go next, is not tested here, and a second demand can also override a preference altogether [10]. Flight selects for a symmetric airframe [12]. A bias that is faint until demanded and cheap when expressed does not contradict that selection. As in humans, whose symmetric arms are used asymmetrically, the asymmetry sits in the use of the body, not in its build. Whether such asymmetries matter ecologically is likely to depend on how often the demanding regime is entered: as thermals become more marginal and winds less predictable under global change [57], the very conditions that elicit the bias could be met more often, and a per-manoeuvre cost that is negligible today could become consequential for the individuals least able to modulate it. Three questions follow: whether demand-elicited laterality scales into fitness under harsher conditions; whether the same latent-then-amplified structure governs other lateralised motor behaviours; and what sets the side in the first place, which nest-tagged cohorts, where siblings share a nest and its rearing conditions, make testable in the wild.

## Statements

### Ethics

The data analysed here were not collected for this study. All tracking data were gathered by established research programmes for their own purposes and under their own permits, and are reused here; the present work involved no fieldwork, no handling of animals and no new deployments. The original approvals are as follows.

#### European honey buzzard (Finland)

Permits to trap and deploy GPS-tracking devices were issued by the Centre for Economic Development, Transport and the Environment in Southwestern Finland (VARELY/8915/2022).

#### European honey buzzard (Switzerland)

Experimental licenses were provided by the Food safety and Veterinary office (LSVW) of the Canton of Fribourg (permit no. N-34775-2022-11-FR) and the Federal Office for the Environment (FOEN) and the Amt für Wald und Natur Freiburg, license number: wNA20240325.1.

#### Golden eagle (Alps)

The handling and ringing of golden eagle nestlings in Switzerland was carried out under the authorisation of the Food Safety and Veterinary Office Grisons, permit nos. GR 2017_06, GR 2018_05E, GR 2019_03E, GR/08/2021, and the Federal Office for the Environment, licence no. TV201903E. In Italy, the permissions for handling, tagging and marking were obtained from the autonomous region of South Tyrol (Dekret 12257/2018 and Dekret 8788/2020), as well as from the region of Lombardia for ringing and tagging through in Lombardia and South Tyrol by ISPRA (Istituto Superiore per la Protezione e la Ricerca Ambientale) with the Richiesta di autorizzazione alla cattura di fauna selvatica per scopi scientifici (l.r. 26/93). In Austria, all procedures for handling, marking and tagging were approved by the Ethics Committee of the University of Vienna (no. 2020-008) and permitted by the Federal Ministry for Education, Science and Research (no. GZ: 2020-0.547.571 & GZ: 2023-0.150.989), Styria (BHLI-165942/2021-2; ABT13-178011/2021-7 & 14; ABT13-106176/2023-6; ABT13-226032/2024-6; ABT13-156808/2025-6), Upper Austria (LFW-2021-263262/20-ÖL), Lower Austria (SBL2-J-1993/002; SBL2-J-1970/006; SBL2-J-1993/003), Tyrol (LZ-NSCH/B-474/3-2023) and Carinthia (10-JAG-2912/2-2023). Finally, in Germany birds handled, tagged and ringed were done so under the permission issued by the government of Oberbayern (2532.Vet_02-16-88 and 2532.Vet_02-20-86). All procedures followed the ASAB/ABS guidelines for the ethical treatment of animals in behavioural research and teaching and all applicable international, national, and/or institutional guidelines for the care and use of animals were followed. The handling of birds was performed with maximum care and minimal disturbance to nests and the landscape.

#### Golden eagle (Sweden)

All procedures for handling and ringing were licensed by the Swedish Museum of Natural History Ringing Centre (Stockholm, Sweden). Tagging was approved by the Malmö/Lund ethical committee (permit 5.8.18-14807/2021).

#### White stork (south-western Germany)

Permits for tagging and tracking the white storks were issued by the authorities of the Federal States (G-13/28 and G-268 15/47 by Regierungspräsidium Freiburg, MPI269 O-1/14 by Regierungspräsidium Tübingen).

### Data accessibility

The data we analysed and the complete analysis code are deposited in the Edmond repository (Max Planck Digital Library): https://doi.org/10.17617/3.X9P38E. The deposit contains the six derived tables the analysis runs on, the full analysis pipeline in R with a runner script, the package versions, a column dictionary, a README, and the results store from which every number in the article and its electronic supplementary material is read. Every result, figure and table in this article and in the electronic supplementary material can be regenerated from the deposited tables and code alone, with no access to the raw data required. The raw GPS and inertial sensor data are archived on Movebank (https://www.movebank.org), subject to the data-use agreements of the programmes that collected them, under study IDs 2201086728 and 2235142320 (European honey buzzard), 282734839 (golden eagle) and 1176017658 (white stork). They are not deposited because they contain nest and territory locations, which are conservation-sensitive and played no part in this analysis.

### Declaration of AI use

We used Claude Code (Anthropic; Claude Opus 4- and 5-series models, 2026) during the preparation of this manuscript. The tool assisted with developing, auditing and documenting the analysis pipeline in R; with verifying that every number reported in the text, figures and tables resolved precisely from the code; and with editing, formatting and L^A^T_E_X mechanics. We carried out the literature search and directed the tool to the sources we had chosen. The tool read each cited work against the statement it is cited for, and it proposed some of the references for the methods we adopted. The tool and the authors both checked every cited reference against its source. The project’s git history documents all changes. The tool was not used to generate data or results, and it did not set the scientific claims. All results derive reproducibly and transparently from the analysis pipeline run on the tracking data. The interpretation, argument and writing of the manuscript are the authors’. We reviewed all AI-generated material in this manuscript, take full responsibility for its content, and confirm that no AI tool is listed as an author.

### Authors’ contributions

K.S. conceived the study. A.F., M.U.G., A.H., P.S., P.B. and E.N. collected the data. E.N., K.S. and E.Y.Y. processed and analysed the data. K.S. and E.N. wrote the first draft of the manuscript. All authors critically contributed to the drafts and gave their final approval for publication.

## Acknowledgements

We are grateful to Jake Graving, Carel van Schaik, Martin Wikelski, Yevgenia Kozorovitskiy, Wolfgang Heidrich, Lochlan Walsh and Alba Motes Rodrigo for their feedback on earlier versions of the manuscript, and to Imran Razik for the illustrations of soaring flight. We are sincerely grateful for the efforts made by Wolfgang Fiedler, Teemu Honkanen, Aki Korhonen, Mikko Honk-iniemi, Ilmari Häkkinen, Janne Leppänen, Hannu Lehtoranta, Inga Kujala, Ilari Soppela, Tomi Hakkari, Helena Siivonen and Kari Ketola for tagging the honey buzzards in Finland. A special thanks goes to Valentijn S. van Bergen for tagging the Swiss Honey buzzards and to Adrian Aebischer, Benedetta Catitti, Frank Haven, Lara B. Gross, and Urs G. Kormann for making the tagging effort possible. We are grateful to Adriano Greco, Alessandro Mercogliano, Andrea Roverselli, Klaus Bliem, Franz Schüttelkopf, Shane Sumasgutner, Varalika Jain, Rebekka Kreikenbohm, Johannes Ploderer and Lucas Landolfi for their support during the eagles’ tagging. We are grateful for the crucial assistance of the mountain rescue teams of the Guardia di Finanza and Grünau/Almtal (Franz Rührlinger) as well as the forestry and game wardens of South Tyrol/Alto Adige and the national park teams in Gesäuse, Kalkalpen, Wildnisgebiet Dürrenstein-Lassingtal and Hohe Tauern, as well as the ERSAF-Stelvio national park Italy and the Department of Wildlife and Fishery Service Grisons, Switzerland and many gamekeepers. We also thank the accompanying veterinarians Drs Michel Mottini and Gilberto Volcan for their time and expertise in the field. We are grateful to Julia Hatzl, David Jenny, Matthias Tschumi, Enrico Bassi for the project organisation and tagging procedures in Switzerland and Italy. We thank Martin Wikelski for supporting the various projects that contributed data to this work. We are deeply thankful for the dedication and hard work by David Folkesson to monitor and ring Golden Eagles in Sweden. Without him tagging would have been impossible.

## Electronic Supplementary Material

### Supplement sections

#### Conventions used throughout

Three values separated by slashes are given in the order honey buzzard / golden eagle / white stork. This is the same order as, and is equivalent to, the labelled triplets of the main text, where the species carrying each number are marked HB (honey buzzard), GE (golden eagle) and WS (white stork). Square-bracketed ranges are 95% confidence intervals unless stated otherwise. A pointer to another section of this supplement is written *(see S8)*. A square-bracketed *[S8]* is a numbered reference in the list at the end. A number in parentheses after an analysis name, as in “the keystone (05)” or “switching cost (35)”, identifies the pipeline element that produced it in the deposited code and the results store. It is an identifier, not a literature reference.

#### Reproducibility

Every number in this supplement and in the main text, and every figure and table, is read from one results store, a single table of estimates, intervals, p-values and sample sizes that the analysis scripts write and nothing else edits. The store is deposited together with the six analysis tables it was computed from, the R pipeline that produces it, the scripts that draw every figure and table, the package manifest and a README that describes the layout and the order in which to run the scripts, in the reproducibility archive named in the Data accessibility statement. Every result reported here regenerates from that archive without access to the raw tracking data. Every analysis is described in words in the text. In the figures, every plotted quantity is the analysis it reports, read from the store or from the saved model objects.

#### Software and statistical tools

All analyses were run in R 4.6.0 [S1]. Tracking data were handled as move2 objects [S2]. GPS tracks were cleaned with the self-thresholding hierarchical outlier cascade of move2utils [S3]. Wind, gust and updraft were estimated from the birds’ own thermalling with moveWindSpeed [S4]. The binomial mixed models of record were fitted in glmmTMB [S5], smooth terms in mgcv [S6], spatial residual diagnostics in spdep [S7] on geometries handled by sf [S8], and cluster-robust covariances with sandwich [S9]. References cited in this supplement are listed at its end and are numbered separately from the main text.

#### Units

GPS was recorded at 1 Hz in bursts (table S1). A circle is one completed net rotation of 360° of the cleaned track. A sustained glide ends the current circle. A circle is valid if its spatial extent is at least 3 m, it is not both under 10 m in extent and slower than 18 degrees per second, its net rotation is at least 300° in magnitude, and its mean ground speed is at least 2 m s*^−^*^1^. The last two exclude circles whose turning sense is ambiguous and stationary GPS jitter. A thermal is a run of valid circles separated by no more than 250 s.

### S1: Sample and cohort accounting

The counts differ between analyses because each quantity estimated requires different coverage, so each analysis includes the birds that carry the data it needs.

The attrition runs from 354 tagged birds to 339 tracked after a-priori exclusions to the analysed cohorts (figure S1a). Every count is derived in code and written to the results store: individuals, circles, thermals, tracked and soaring bird-days, IMU-recorded circles and VeDBA windows (vectorial dynamic body acceleration, the accelerometer measure of effort defined in S4), per species and per stage, with the per-individual distributions (circles, hard circles and hard circles per bird). table S1 gives the study-system summary.

**figure S1:**
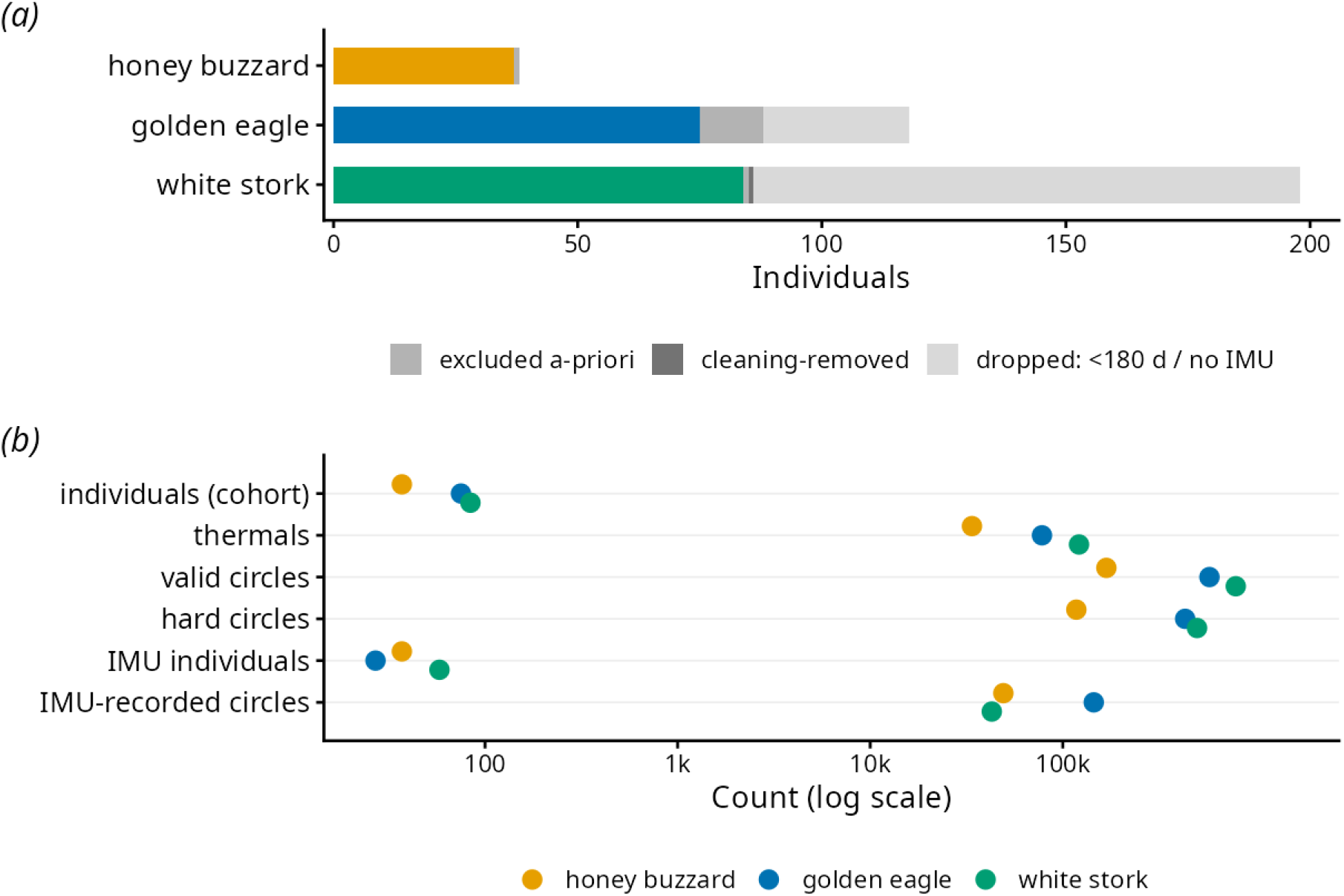
Sample and cohort accounting. (*a*) Attrition from tagged birds to the analysed cohort, per species. From 354 tagged birds, 339 remained after a-priori exclusions (monitoring or rehabilitation animals, wrong-population and tag-swap cases), and 196 entered the final cohort after the post-cleaning longitudinal filter (at least 180 days of GPS, or IMU circles): 37 honey buzzards, 75 golden eagles and 84 white storks. The final cohort is drawn in the species colour and the removed birds in grey. (*b*) Sample size at every analytical level, per species, on a logarithmic scale: individuals, thermals, valid circles, hard (above-onset) circles, IMU-instrumented individuals and IMU-recorded circles. The counts are given in table S1, except the hard circles, 117 595, 431 905 and 497 904. Inclusion is set by what each quantity requires, giving three analysis sets: the difficulty analyses (37 / 76 / 135 = 248 birds), the longitudinal cohort (37 / 75 / 84 = 196) and the IMU set (37 / 27 / 58 = 122).

**Table S1.** Study systems. Counts, geographic ranges and year spans are code-derived. Tagging region and sensor sampling are study-configuration constants (Methods, S2). Circles, thermals, acts and bird-days are the totals for the 196-bird descriptive cohort defined below.

| Attribute | European honey |  |  |
| --- | --- | --- | --- |
|  | buzzard | Golden eagle | White stork |
| Tagging region | Finland (31) and<br>Switzerland (6) | European Alps and<br>southern Sweden | south-western<br>Germany |
| Tagged individuals | 38 | 118 | 198 |
|  | buzzard | Golden eagle | White stork |
| Analysed cohort<br>(IMU or $\geq 180$ d<br>GPS) | 37 | 75 | 84 |
| IMU-instrumented<br>individuals | 37 | 27 | 58 |
| Birds with valid<br>circling data | 37 | 76 | 135 |
| Classified lateralised<br>/ symmetric ($\geq 100$<br>hard circles) | 29 / 8 | 53 / 18 | 85 / 33 |
| Deployment years | 2022–2024 | 2017–2025 | 2020–2026 |
| Data years | 2022–2026 | 2018–2026 | 2020–2026 |
| Longitude range<br>(degrees) | −14.4 to 49.6 | 6.2 to 17.4 | −16.5 to 10.9 |
| Latitude range<br>(degrees) | −27.0 to 63.3 | 44.8 to 62.4 | 13.5 to 51.8 |
| Sensor types on the<br>tag | 4 | 4 | 4 |
| GPS sampling | 1 Hz (in bursts) | 1 Hz (in bursts) | 1 Hz (in bursts) |
| IMU orientation<br>sampling | 20 Hz (quaternion) | 20 Hz (quaternion) | 20 Hz (quaternion) |
| Valid thermalling<br>circles | 168 398 | 577 440 | 790 981 |
| Thermals | 33 760 | 77 975 | 121 406 |
| IMU-recorded circles<br>(at least a half-turn) | 49 132 | 144 825 | 42 793 |
| IMU bursts | 374 350 | 134 923 | 775 655 |
| Tracked bird-days | 15 082 | 65 228 | 65 309 |
| Soaring bird-days | 3 758 | 13 544 | 14 369 |

Inclusion is set per analysis family. The difficulty analyses (the per-individual odds model and its repeatability, development and robustness checks) use every bird with sufficient valid circles, with per-bird classification requiring at least 100 hard circles: N = 37 / 76 / 135 (248 birds). The relationship is within-circle and has no longitudinal requirement, so all data are retained for maximal power. The descriptive cohort and the population-level existence test use birds with valid GPS circling data that also contributed IMU circles or at least 180 days of GPS: N = 37 / 75 / 84 (196 birds), because characterising a bird as a persistent unit across the annual cycle requires longitudinal coverage. The IMU cost analyses (the per-manoeuvre instability, climb, load-factor, switching and daily-effort analyses) use the instrumented set: N = 37 / 27 / 58 (122 birds). It is not a subset of the longitudinal cohort: 25 white storks with IMU circles had no usable GPS circling and enter the IMU analyses only. Life-cycle staging uses the same longitudinal cohort.

### S2: What the IMU can and cannot deliver

The laterality analysis uses orientation-invariant quantities from the tag, which survive an unknown mount angle. The absolute bank angle does not, and is not used.

Two distinct effects are both loosely called tilt, and we keep them apart throughout. *Mount tilt* is the fixed rotation between the tag and the bird, which is removable as a per-bird offset. Quantities from which it has been removed are called mount-offset-corrected. *Acceleration tilt* is the displacement of the apparent-gravity vector by centripetal acceleration during a sustained turn, which is not removable and is why absolute bank fails: gyroscopes measure angular velocity but accelerometers measure specific force, so during dynamic motion the accelerometer output cannot be read as the gravity vector, and an attitude estimate that uses it as one underestimates the bank, or in the ideal coordinated turn cannot identify it at all. A quantity that is immune to both, because it is differenced, we call *orientation-invariant*.

Raw accelerometer and quaternion streams are converted to Euler roll, pitch and yaw, then filtered and handled per burst. The manufacturer’s documentation is not publicly available, so we state the conversion here and this supplement is self-contained. The tag returns the quaternion as four 16-bit integers 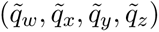. The scalar part is scaled directly, while the vector part is rescaled so that the quaternion has unit norm,

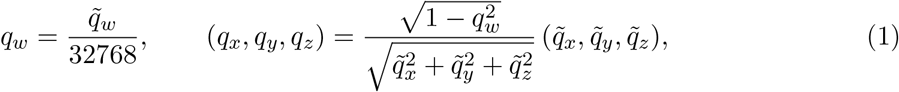

with the vector part set to zero where 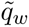 = *−*32768 (scalar *±*1). The Euler angles in the tag frame then follow as

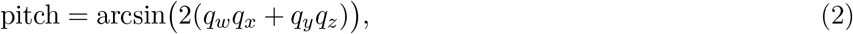

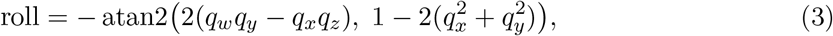

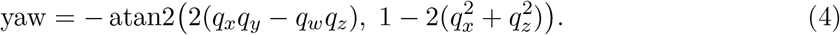

The laterality estimate uses only differenced yaw and differenced roll, both of which are invariant to a constant mount offset. The global distributions per species (figure S3) carry a sanity check: absolute roll approaches 180° (an apparent inversion) in fewer than 0.05 per cent of samples in every species, and pitch exceeds 60° in about a tenth of honey-buzzard samples and almost never in the other two. Where they do, the cause is the apparent-gravity corruption of attitude during sustained circling, not genuine acrobatics.

higher on the non-preferred side (from preferred to non-preferred: 13.2→13.7, 10.2→10.9, 13.0→13.1 degrees per second), and the instability asymmetry survives the control in the two raptors (the non-preferred side still carries 1.2% more instability, p = 0.008, in the honey buzzard and 0.9%, p = 0.032, in the golden eagle, down from 2.2% and 2.9% without the control but still significant). Conditioning on a variability that may share a cause with in-flight instability is if anything over-conservative, so this survival confirms the asymmetry is not an artefact of more-variable banking on the weak side. The absolute side differences in roll SD, 0.045° and 0.059° (honey buzzard and golden eagle), exceed even the e-obs single-sample resolution (∼0.011° per least-significant bit, 15-bit), a conservative floor since each instability value already averages over thousands of samples. The white stork remains equivocal (a borderline conditioned p = 0.048 on a negligible −0.002° difference; its primary effect is null, p = 0.23).

Two provenance checks close the dossier. The tag mount offset changes little over a deployment: fitting the level-cruise reference roll on date for every tag with enough reference bursts gives a median absolute change over the whole deployment of 2.9, 3.0 and 5.0 degrees (111 of 122 tags), That corruption is quantified directly. On the GPS-IMU overlap the mount-offset-corrected absolute IMU bank reads only about 6 to 7° against a kinematic bank of about 23°, a slope near 0.2 (figure S2): the apparent-gravity lock that a gravity-referenced attitude estimate is expected to show whenever external acceleration is sustained, here reproduced on free-flying birds. We therefore do not use absolute bank anywhere in the analysis.

**figure S2:**
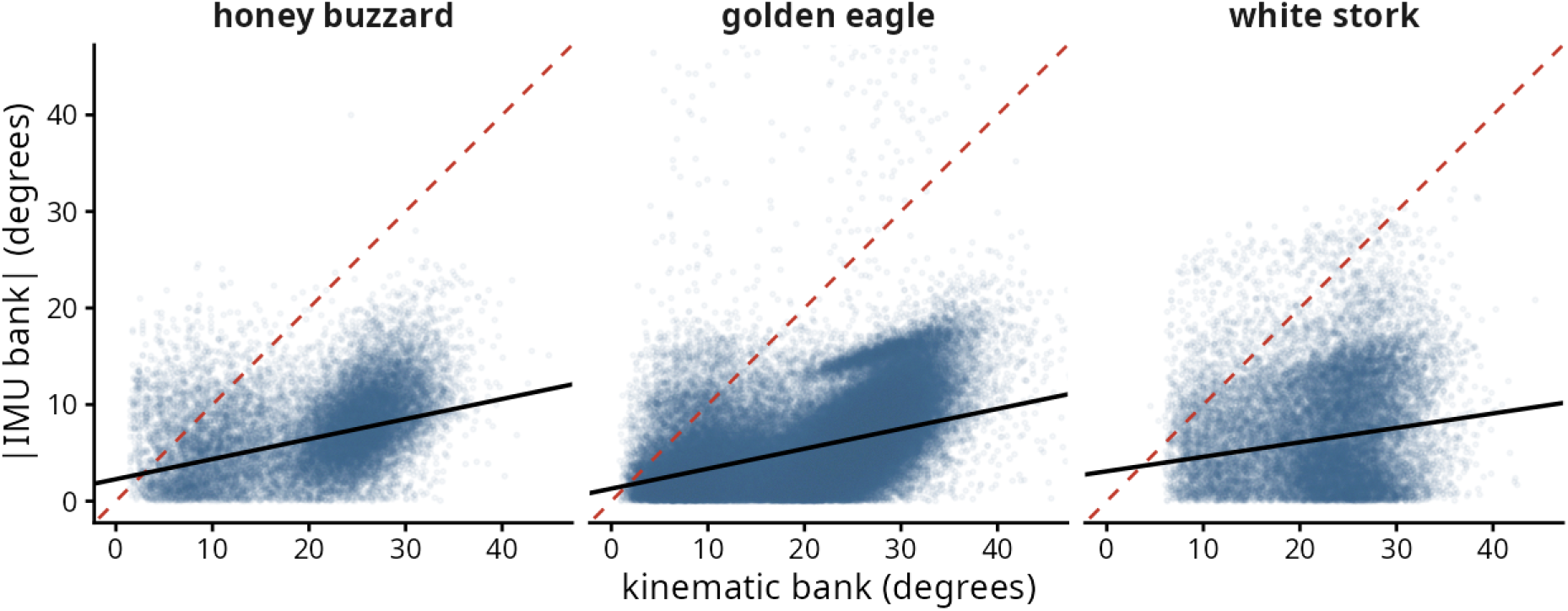
Absolute IMU bank cannot be trusted, so bank is taken kinematically. On the GPS-to-IMU overlap, per species, the mount-offset-corrected absolute IMU bank (the e-obs quaternion roll minus a per-bird mount offset) against the kinematic bank the bird must hold to fly the circle. The physically required bank is about 23 degrees, but the mount-offset-corrected IMU registers only about 6 to 7 degrees (a magnitude shortfall of 14 to 17 degrees; the bird-level tracking slope, the black line in each panel against the dashed line of equality, is 0.21, 0.21 and 0.15), reproducing on our own birds the apparent-gravity lock that corrupts absolute roll precisely while circling. The direction of the lean survives well in the two raptors (bank-direction consistency 0.91 and 0.86) but only weakly in the white stork (0.48), whose larger and more variable mount offset makes even the direction of its absolute IMU bank less reliable. This does not affect the quantity estimated, which reads laterality direction from GPS and the orientation-invariant yaw (where the stork bridge agreement is 99.8 per cent), but it is why IMU-derived stork quantities carry lower confidence, consistent with the stork instability asymmetry itself being null. Its magnitude does not survive in any species, which is why we take bank kinematically from the track and let the IMU contribute only orientation-invariant differenced quantities.

The instability metric needs no absolute bank. Wobble is the short-window **differenced** roll SD, not absolute roll, and differencing cancels both the constant mount offset and the slowly varying apparent-gravity component, so the metric is orientation-invariant by construction, for the same reason the laterality estimate reads GPS and yaw direction rather than absolute bank. The side comparison is additionally conditioned on turn rate at matched tightness, removing the mean relationship between turn rate and in-flight instability.

#### Resolution

The side difference in in-flight instability is small in degrees (0.045 / 0.059 / −0.002° of within-second roll SD on a per-bird median of 2.3 / 3.0 / 2.1°) and is not resolved by any single sample: it is the within-bird mean over 49 132 / 144 825 / 42 793 paired IMU-recorded circles, and its test is the per-bird paired test reported in the main text. The sensor noise floor is the same on both sides of the contrast, being the same tag on the same bird, and cancels in the pairing. The single-sample resolution given below (about 0.011° per least-significant bit) is an order of magnitude below the difference.

One residual worry is that matching mean turn rate does not match the variability of banking within a circle, which could inject apparent-gravity variance into the differenced roll. Adding the within-circle SD of per-second net yaw (an orientation-invariant proxy for within-circle bank variance) as a covariate and refitting shows the confound is real but small: the within-circle net-yaw SD is slightly and laterality is in any case a direction, invariant to a fixed or slowly changing offset (figure S3b). And one golden eagle tag (e-obs 7503) with a faulty orientation sensor is excluded from every IMU analysis while its GPS is retained, since the two arms are independent.

**figure S3:**
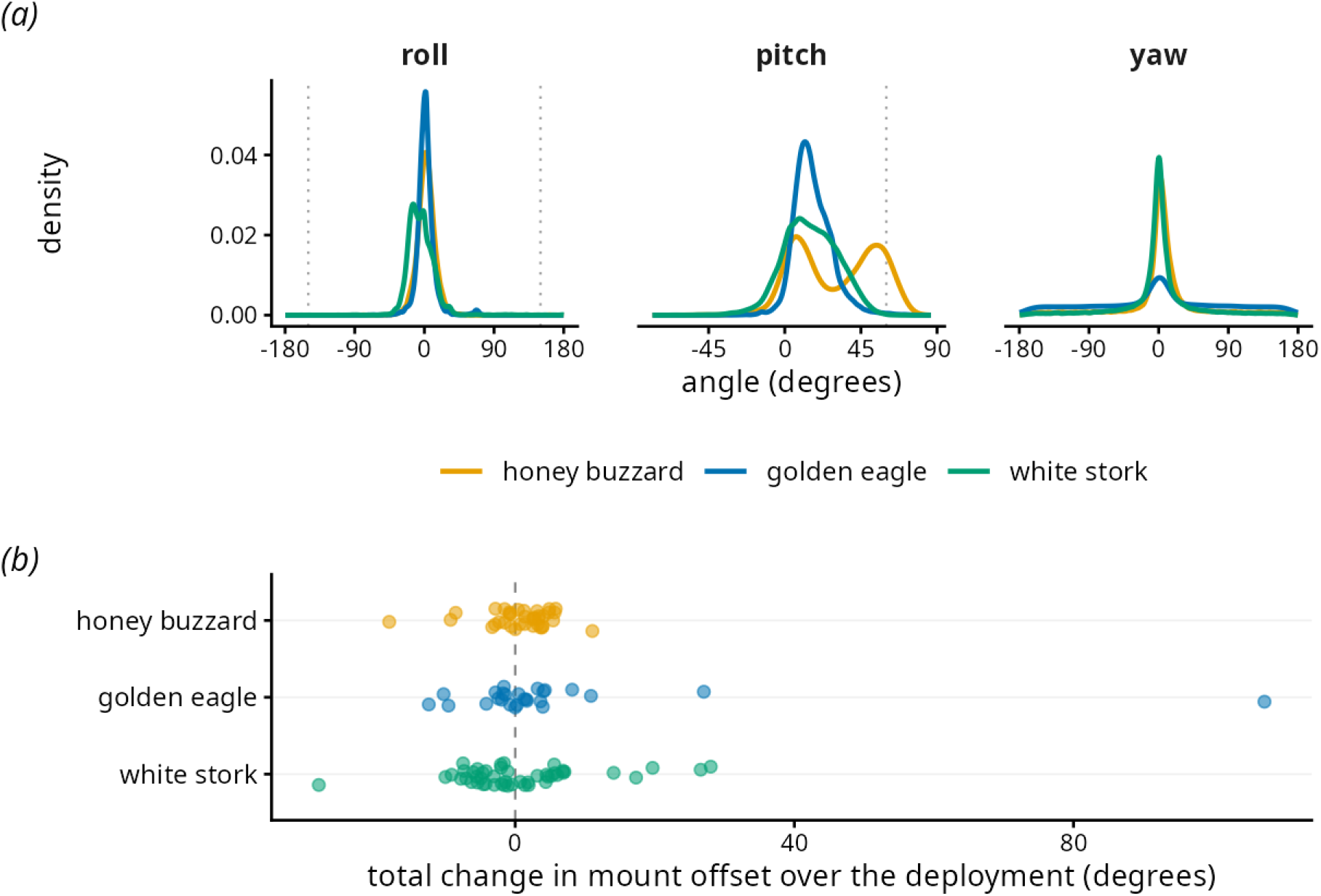
The IMU orientation dossier. (*a*) Global distributions of the derived roll, pitch and yaw angles per species, from the decoded per-sample quaternions. The dotted lines mark the two bounds checked. Absolute roll rarely approaches 180 degrees (|roll| > 150 degrees in 0.05 per cent or fewer of samples in every species), so the tags do not record birds flying inverted. Pitch exceeds 60 degrees in 10.9 per cent of honey-buzzard samples but essentially never in the eagle or stork, the signature of the apparent-gravity corruption that inflates the reported attitude during sustained circling. (*b*) The change in tag mount offset over each deployment: for every tag with at least ten level-cruise reference bursts spanning at least thirty days, the per-tag slope of the reference roll on date times that tag’s own deployment span. The median absolute change is 2.9, 3.0 and 5.0 degrees over 37, 26 and 48 tags. One golden-eagle tag changed by 107 degrees.

### S3: The GPS-IMU bridge

The turning direction is read from the GPS track, while the per-manoeuvre cost signals are read from the IMU, so the two instruments must agree on which way each circle turned.

On circles seen by both sensors, the turning direction from GPS track curvature and from the orientation-invariant IMU yaw agree on 99.0% of circles in the honey buzzard, 98.4% in the golden eagle, and 99.8% in the white stork.

### S4: The quantity estimated and the models

The claim that a turning-side bias is an individual trait rather than a population handedness is a statement about the between-individual variance in the per-bird side. Reducing each bird to a raw laterality index [S10] and thresholding it discards the estimate’s uncertainty and treats a bird with fifty circles like one with fifty thousand [S11].

#### Definitions

The response of the main model is *R*, whether a circle was flown rightward (1) or leftward (0). The response of every conditioning model is *toward*, whether a circle was flown to the bird’s preferred side, where the preferred side is the bird’s majority side over all its circles, easy and hard alike. It is never assigned on hard circles alone, so the gain is not built in by the side definition, and the floor is validated on circles the side rule did not use (see S5). A circle is *hard* when its turn rate is at or above the species onset (27 / 15 / 21 degrees per second) and *easy* below it. The difficulty term is the hinge *g_z_* = max(0*, ω −* onset)*/*sd(*ω*), so every gain odds ratio is per one standard deviation of turn rate above the onset. That standard deviation is 12.7 / 9.9 / 8.7 degrees per second over all birds’ circles (12.7 / 9.9 / 8.7 for the longitudinal cohort). An IMU-recorded circle (a *directed act* in the code) is a net rotation of at least 180° accumulated within one burst. *In-flight instability*, written wobble in the model formulas of table S2, is the standard deviation of roll within each second of an act, averaged over the act. VeDBA is computed per 8 s accelerometer window after removing the static component over a 1 s window. In the daily models the daily count of non-preferred circling (the *dose* in the code) is the sum over a bird-day of its non-preferred circles, each weighted by its difficulty, with the preferred-side count entered alongside as a second count. It is never a proportion. Residuals are Bernoulli at the circle, so overdispersion is not identifiable at the observation level. Extra-binomial clustering is absorbed by the thermal and individual random effects, and its residual trace is what the lag-1 and Moran checks measure (see S7).

The main model is a binomial GLMM on the raw turning side of every circle, with nested individual and thermal random effects and the floor-plus-gain hinge on difficulty, fitted in glmmTMB. The analysis is frequentist and stated as such: intervals are profile or Wald confidence intervals throughout. Per-individual shrinkage is intrinsic to the random effect, and the between-individual variance is the object of inference.

That object makes the trait claim precise. “A side-bias is an individual trait” is the between-individual SD of the baseline preference being greater than zero: 0.20 / 0.18 / 0.21 on the log-odds scale, with profile intervals excluding zero. It is not the population mean probability of a rightward circle, which sits at parity by construction and is a footnote. The bias is a graded spread of preferences around parity (antisymmetry *sensu* Palmer 1996), so “individual, not population” means this variance is positive, not that birds split into two discrete handedness morphs, a stronger claim we do not make.

A low circle-to-circle repeatability of the side (ICC 0.013 / 0.013 / 0.026 [S12]) might look like weak consistency, but it is low by construction. A one-bit outcome carries an irreducible latent residual, and because laterality is elicited by difficulty, with the floor near parity on easy circles, an ICC pooling easy and hard circles is diluted by design. A low thermal-to-thermal ICC is what a *conditional* trait must look like. Measured where the trait is expressed, on hard circles, consistency is high: the between-bird SD rises to 0.37 / 0.30 / 0.44, a bird’s early and late preferred-side probabilities correlate at 0.62 / 0.65 / 0.59, and on days on which a bird visibly favoured one side when circling was hard, birds flip side less than chance (permutation p 0.002 / 0.019 / 0.001). Repeatability rises monotonically from the easy to the hard regime, as a demand-elicited trait predicts. This is also why we do not pre-compute a thresholded index (a worked demonstration shows thresholding mis-ranks birds of unequal N), and why any “per cent lateralised” statement is given as a full threshold-sensitivity curve rather than an ad-hoc cut (figure S4).

**figure S4:**
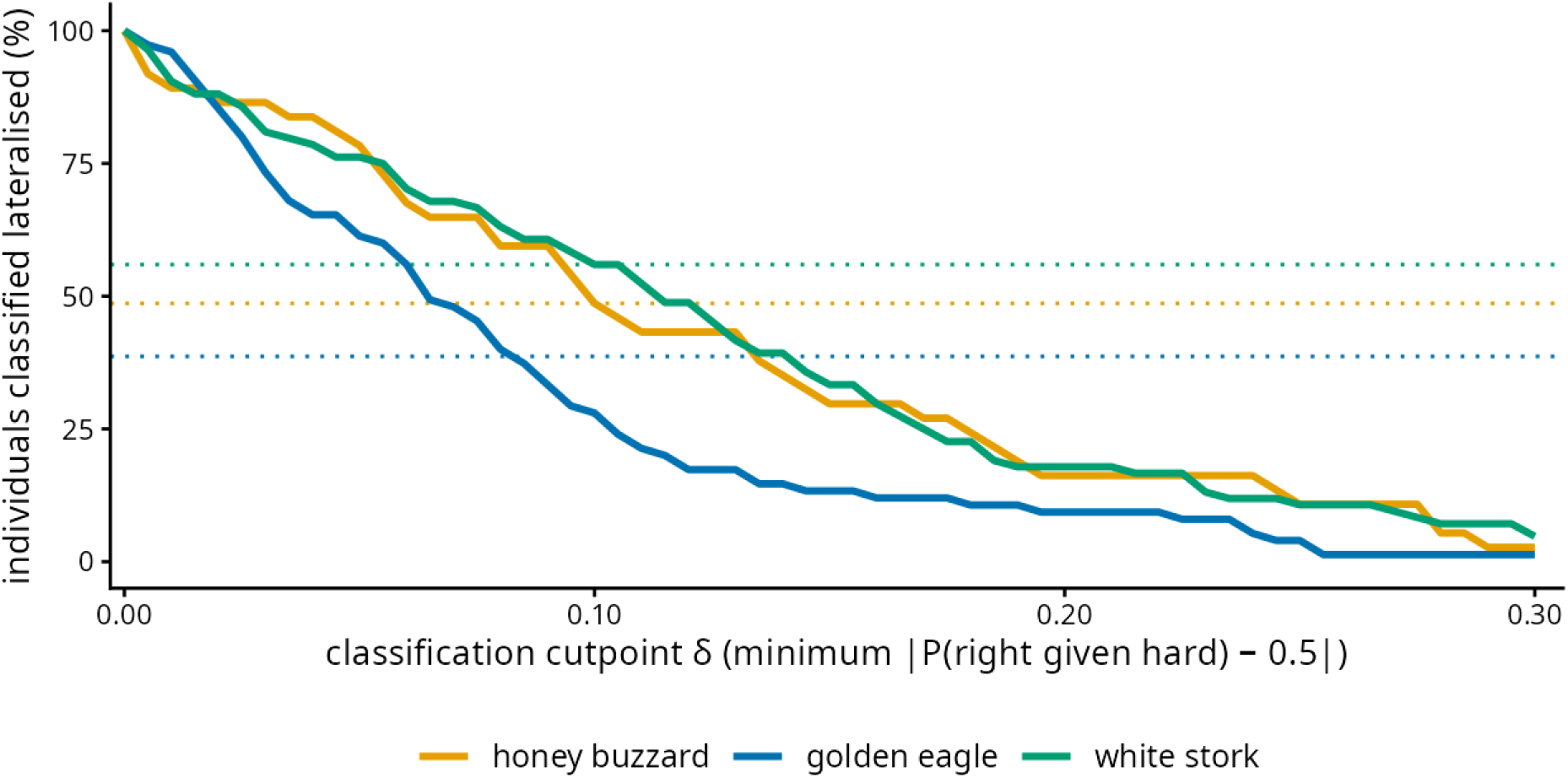
A single “per cent lateralised” depends on an arbitrary cutpoint. The percentage of individuals classified as lateralised, as a function of the deviation-from-parity cutpoint applied to each bird’s fitted probability of a rightward hard circle (solid lines). It falls smoothly as the cutpoint rises, with no natural break, so any single “per cent lateralised” figure is meaningless without stating the cut. The sample-size-aware alternative, the fraction of birds whose 95 per cent confidence interval excludes parity (dotted), does not depend on a deviation cutpoint and is 49, 39 and 56 per cent in honey buzzards, golden eagles and white storks.

On hard circles a bird turned to its own side on 0.55 / 0.55 / 0.56 of occasions, and on the tightest tenth of its circles on 0.58 / 0.57 / 0.60. On those tightest circles 19 / 11 / 17 per cent of birds reached at least a 70:30 split, while over hard circles as a whole almost none did (0 / 0 / 2 per cent). Counted over days rather than circles, on the days a bird favoured one side while circling hard, that side was its own on 0.70 / 0.67 / 0.72 of days.

Full model cards (formula, family, random effects, difficulty parameterisation, N, key coefficients with intervals, residual checks and the convergence record) are given in table S2.

**Table S2.**
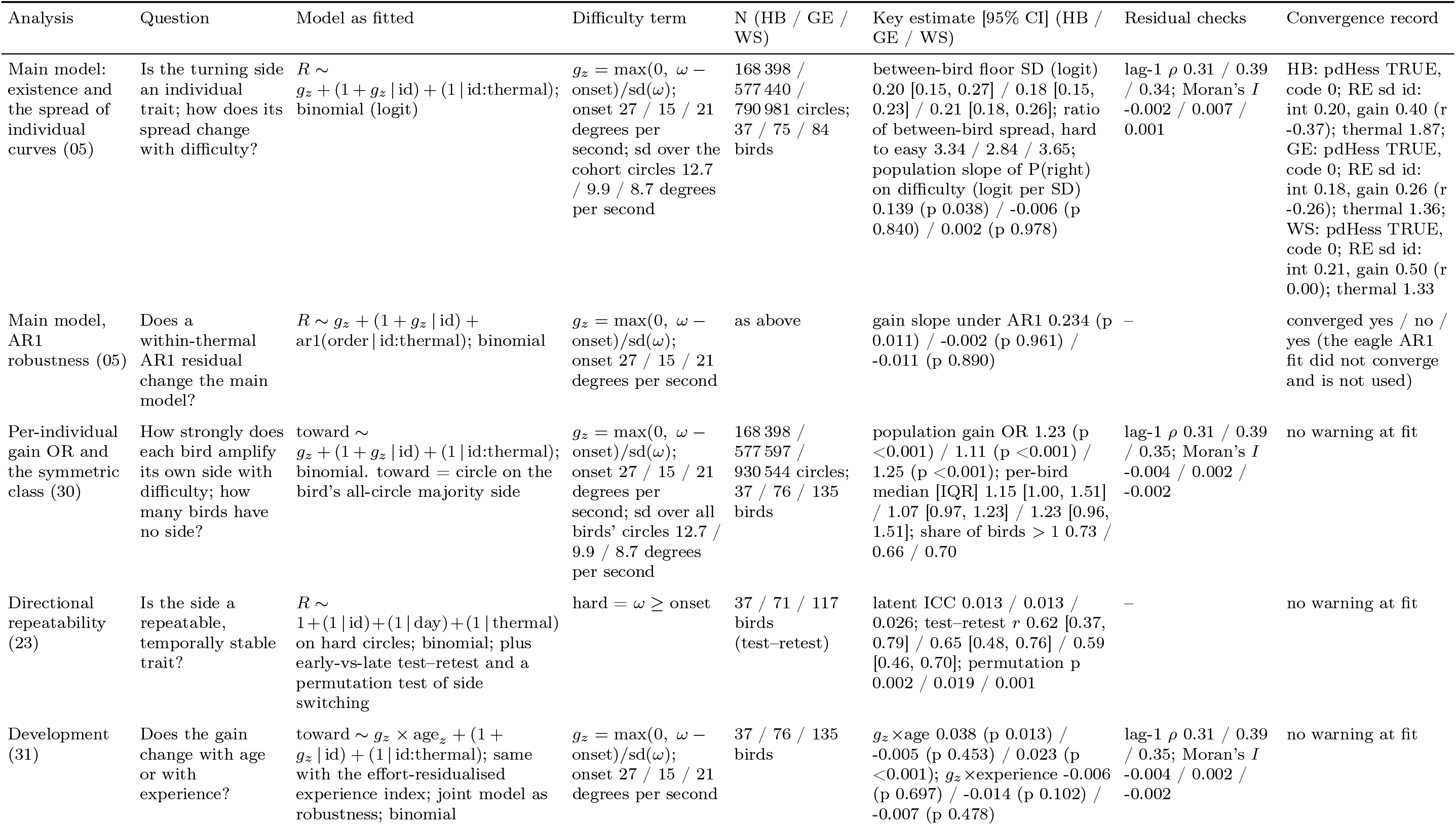
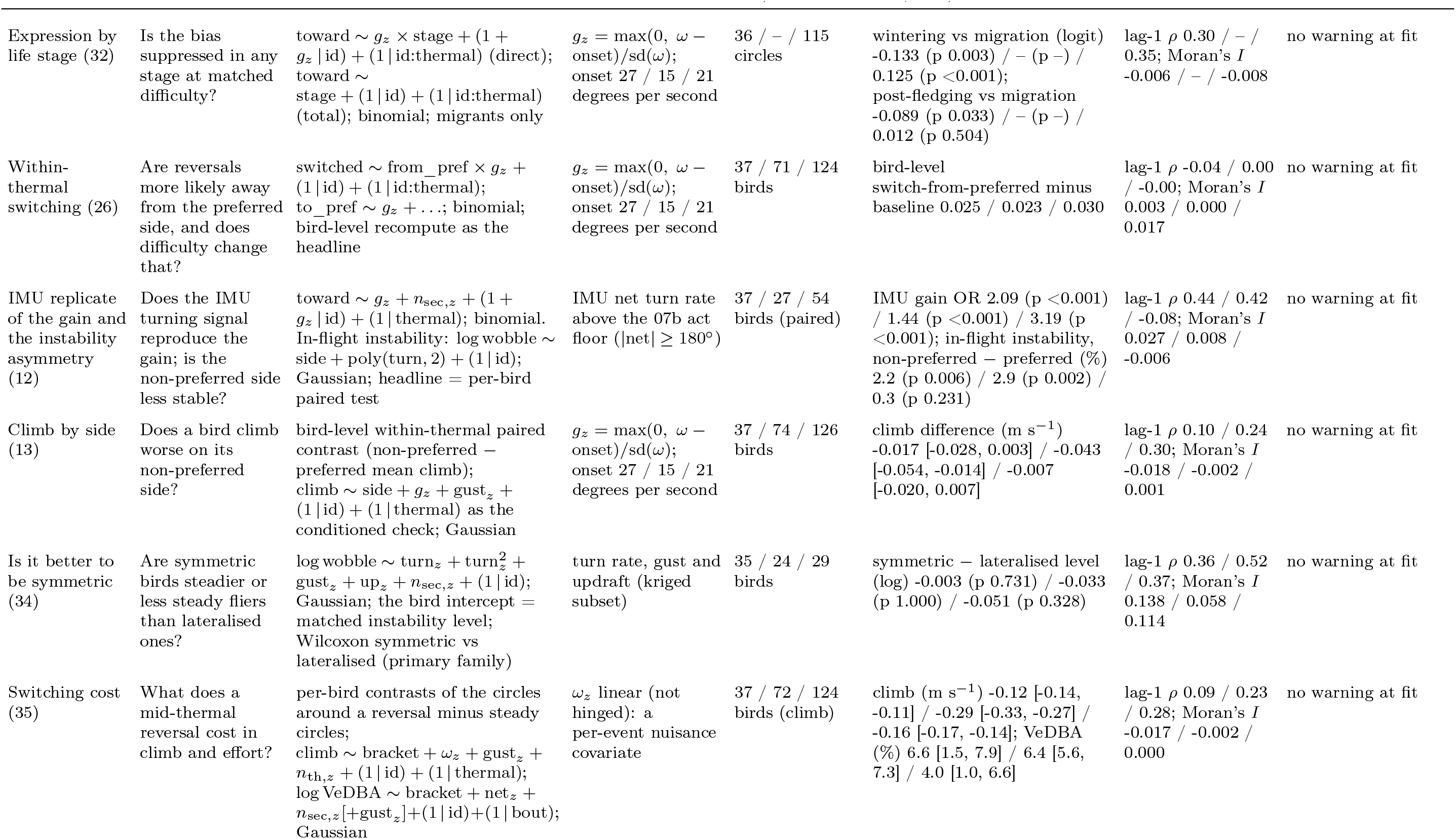
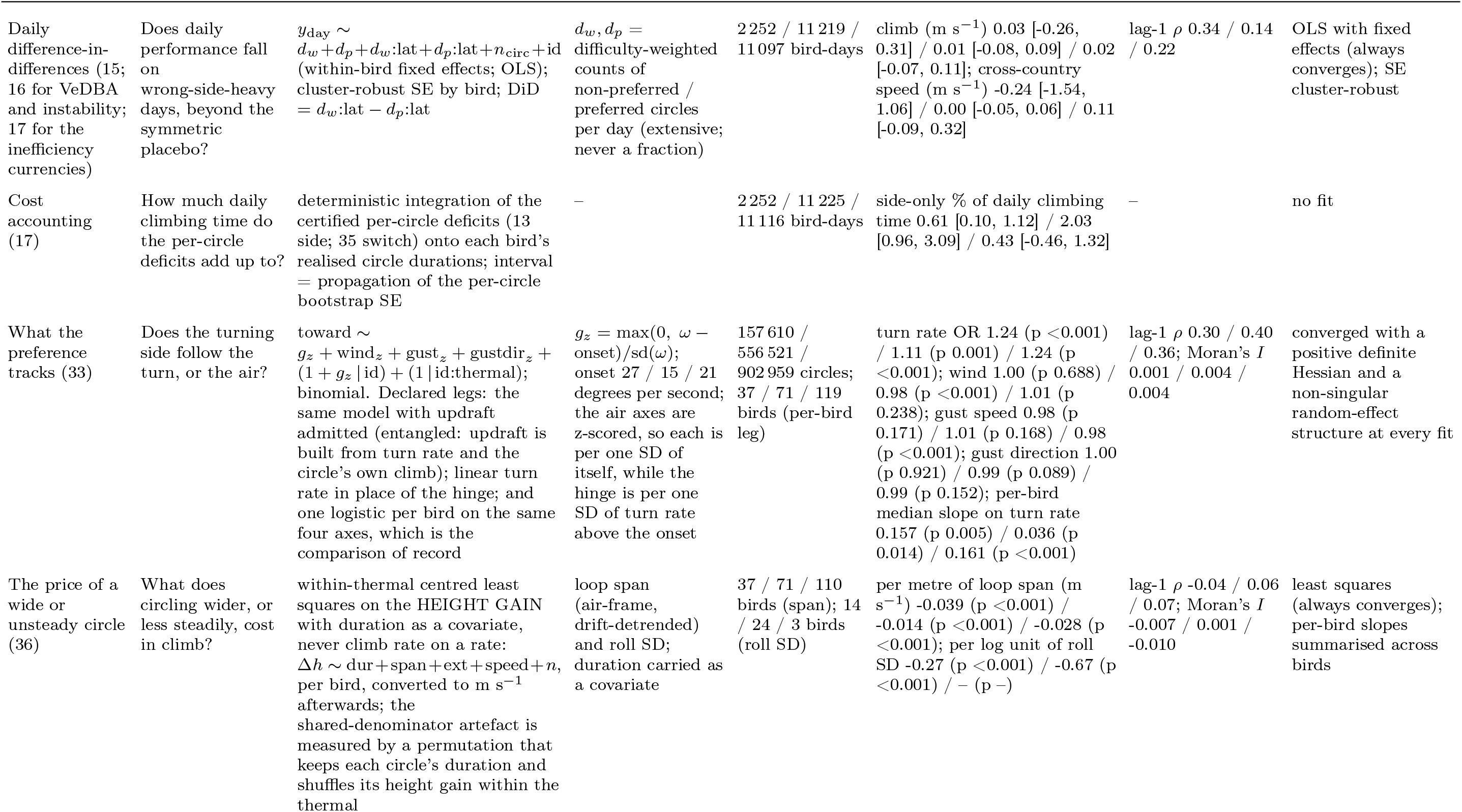
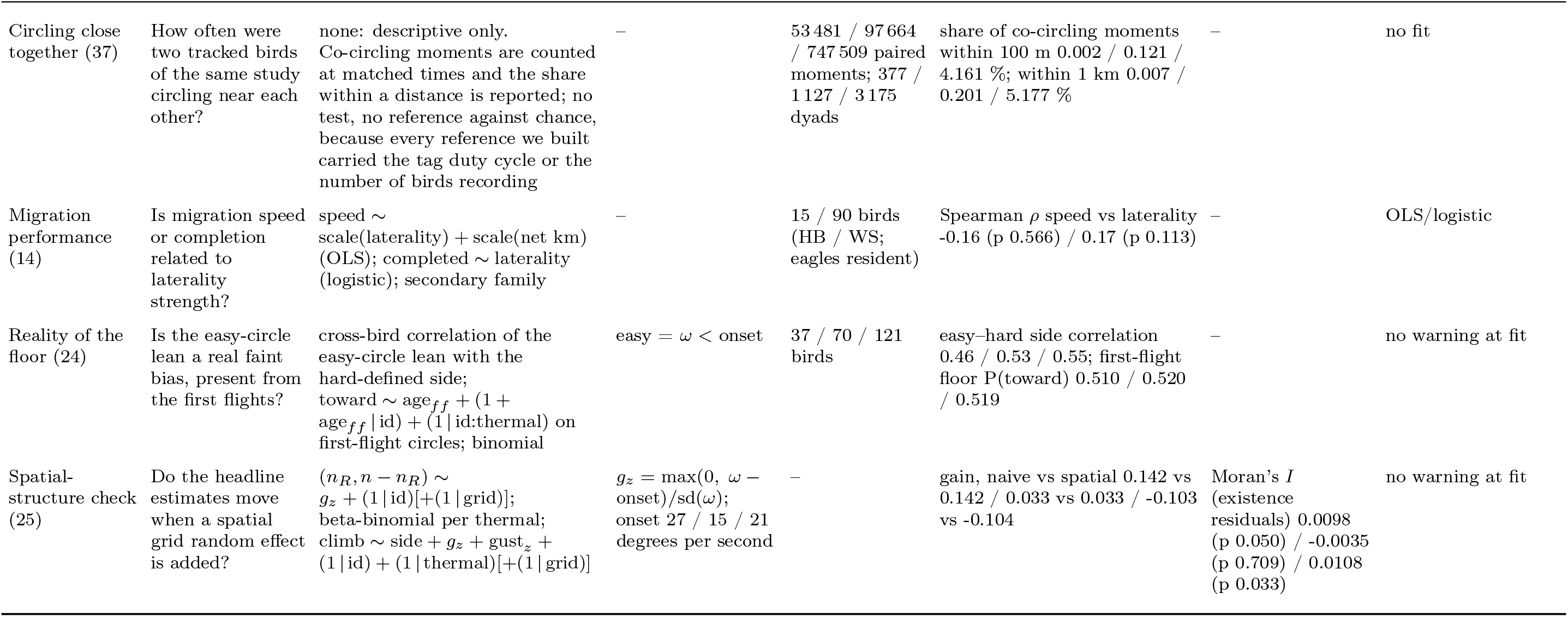
Model cards. One row per model: the question it answers, the model exactly as fitted (response *∼* fixed effects + random effects; family), the difficulty term, the sample behind it, the key estimate, the residual-independence checks and the convergence record. *R* = rightward circle (1/0); toward = circle flown to the bird’s all-circle majority side; *ω* = circle turn rate (degrees per second); *g_z_* = the floor-plus-gain hinge. Subscript *z* marks a standardised covariate: *n*_sec_*_,z_* the seconds recorded in an act, *n*_th_*_,z_* the circles in a thermal, *n*_circ_ the circles in a day, up*_z_* updraft, net*_z_* net rotation, gust*_z_* gustiness, age*_ff_* days since first flight; bracket = a circle adjacent to a reversal, bout = a continuous recording burst, from_pref / to_pref = leaving / joining the preferred side, lat = lateralised versus symmetric, grid = a spatial cell, order = position within a thermal. All glmmTMB fits use the Laplace approximation with nested random effects. Species order honey buzzard / golden eagle / white stork throughout. Intervals on the bird-level contrasts (13, 35, 17) are bootstrap percentiles and need not be symmetric about the estimate. Every number is read from the results store or the saved main-model objects.

#### Symmetric birds

Each bird with at least 100 hard circles (turn rate at or above the species onset) was classified from the Wilson 95% interval of its proportion of rightward hard circles: lateralised if the interval excluded 0.5 and symmetric if it included 0.5 (with at least 100 hard circles the interval is always narrower than 0.20). Lateralised / symmetric: 29 / 8 honey buzzards, 53 / 18 golden eagles, 85 / 33 white storks. At a per-bird 95% interval about one truly symmetric bird in twenty is expected to be classed lateralised, one to two birds per species. The classification is made where the bias is expressed, on hard circles. Figures S4 and S5 count the birds whose fitted probability of turning right at hard difficulty excludes parity (49 / 39 / 56 per cent), and the two differ because the floor is faint by construction and because the counts rest on different denominators, 37 / 71 / 118 birds with at least 100 hard circles against the 37 / 75 / 84 of the longitudinal cohort. Lateralised in the main text means the hard-circle class. The classification is descriptive. Every inferential statement about laterality uses the continuous per-individual odds, and the symmetric birds are the low end of that continuum, not a separate category. No inference depends on the interval width: the class enters only as the placebo arm of the daily difference-in-differences (section S8), where a wider or narrower symmetric class changes the placebo’s composition but not the quantity estimated, and in the between-bird stability comparison (section S8).

**figure S5:**
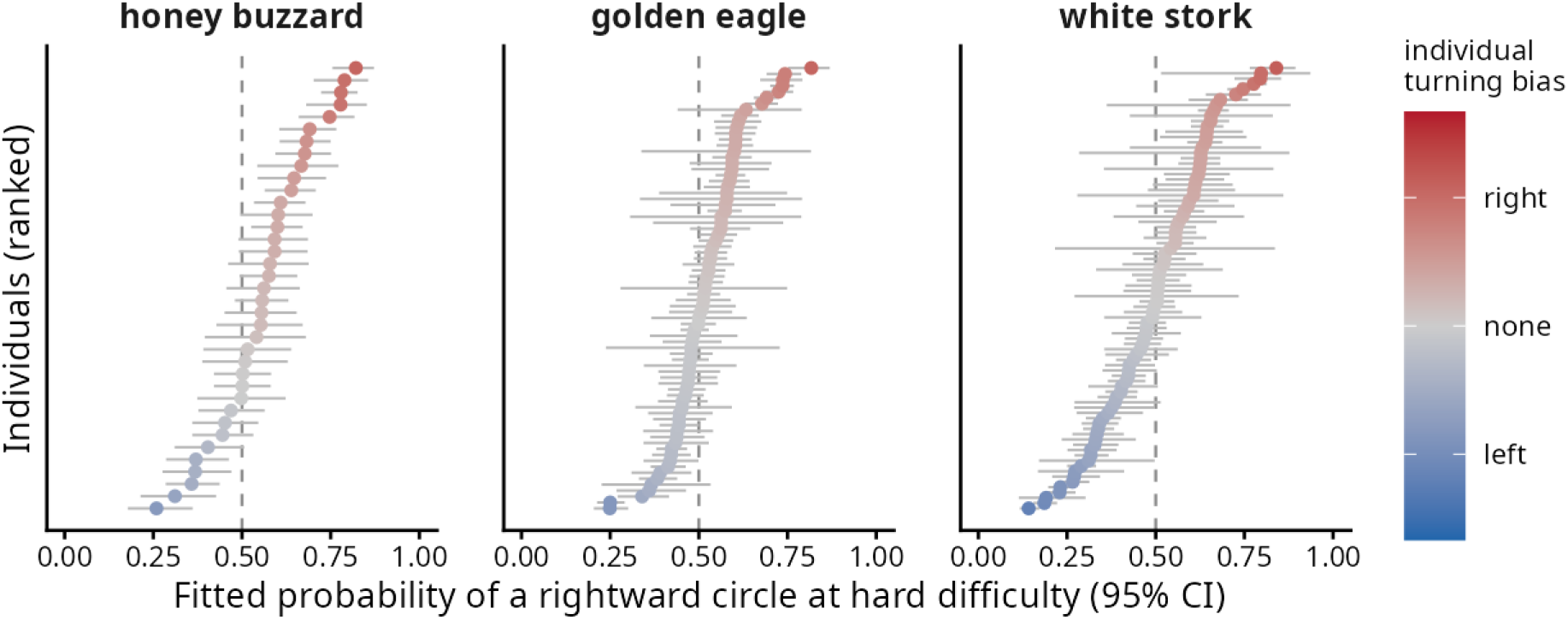
Per-individual coefficients. Each bird’s fitted probability of a rightward circle at high difficulty, with its 95 per cent confidence interval, from the floor-plus-gain main model, one row per bird, sorted, and coloured by turning bias on the same diverging scale as main-text Figure 2. The population sits at parity (dashed line) while individuals fan to their own sides. The between-individual standard deviation of the baseline preference is 0.20, 0.18 and 0.21 on the log-odds scale, and 18, 29 and 47 birds have a confidence interval that excludes parity, in honey buzzards, golden eagles and white storks.

### S5: Difficulty, the onset, and the reality of the floor

Two things could undermine the amplification result: the difficulty threshold (the onset) might have been chosen to produce it, and the faint easy-circle bias (the floor) might be an artefact rather than signal.

The difficulty axis is the circle’s turn rate, which maps to the kinematic bank of a coordinated turn (Methods). The species onset (27 / 15 / 21 degrees per second) is located once, uniformly for every species and outcome, by grid search plus profile-likelihood, and its warrant is the hinge-versus-linear model comparison, which favours the hinge by ΔAIC 87 / 17 / 483, not a p-value tuned on the same data. Because the onset is fixed by this pre-specified procedure and not re-chosen per analysis, it closes the researcher-degrees-of-freedom concern.

The flagship result does not depend on where the onset sits. Refitting the exact headline model across a grid of onsets spanning each species’ ΔAIC-95% CI plus a *±*6 degrees per second margin leaves the population gain OR essentially flat within the CI band (1.22 to 1.23 in the buzzard, 1.09 to 1.11 in the eagle, the weakest case, and 1.25 in the stork), and above one across the entire margin in all three species. At the chosen onset it reproduces the headline odds ratios exactly (1.234 / 1.108 / 1.246). The onset is therefore a descriptive marker, and the amplification is not an artefact of knee placement. (figure S6)

**Figure S6:**
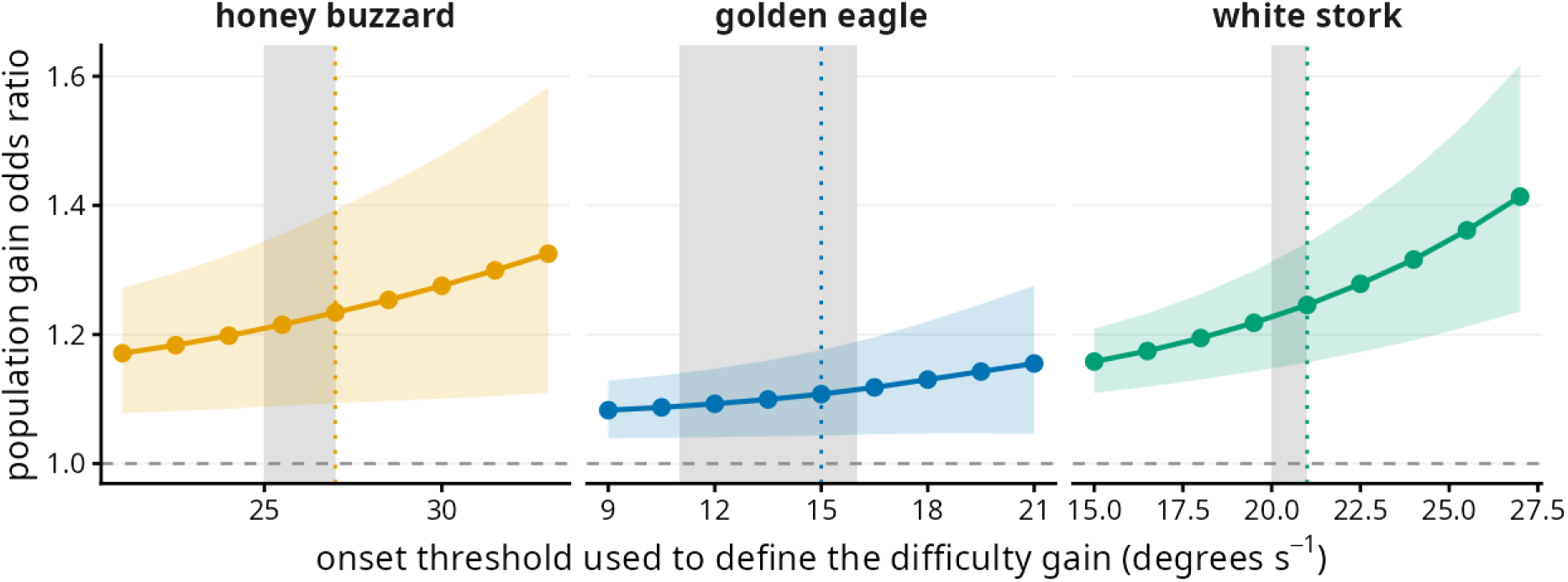
The flagship gain odds ratio stays above one across the onset grid. The population per-individual gain odds ratio, exp(gain slope) from the headline model, refit across a grid of onset thresholds spanning each species’ profile-likelihood ΔAIC-95% confidence interval plus a *±*6 degrees per second margin (points, with 95 per cent Wald intervals as the coloured band). The dotted line is the data-derived chosen onset and the grey vertical band is its ΔAIC-95% CI. Within that band the odds ratio is essentially flat (about 1.22–1.23 in honey buzzards, 1.09–1.11 in golden eagles, the case with the weakest onset evidence, and 1.25 in white storks), and it remains above one across the entire margin in all three species. Refitting at the chosen onset reproduces the headline odds ratios exactly. The amplification is therefore not an artefact of where the difficulty knee is placed (section S5).

The floor is real on three independent counts: each bird’s easy-circle lean aligns with its own hard side (across-bird correlation +0.46 / 0.53 / 0.55), it is already present at a bird’s first recorded flights (the easy-circle probability of turning to the preferred side is 0.510 / 0.520 / 0.519, already above one-half) and flat with age, so only the gain develops, and eagle-sibling similarity is null and underpowered (siblings share a side in 4 of 7 pairs, sibling-identity ICC 0, permutation p = 0.52), placing any innate component beyond the reach of telemetry. Nor is the floor or the spread of individual curves an artefact of selective wind-fit failure thinning the easy end: wind-fit failure is rare and not concentrated on easy circles, and requiring a successful wind fit leaves the spread essentially unchanged and reproduces the floor. Stage comes from an independent latitude state machine (see S6) and does not enter the difficulty axis.

#### What the preference tracks

Tighter circles raised the odds that a bird flew to its preferred side: odds ratio 1.24, 1.11 and 1.24 per standard deviation of turn rate. Wind speed, gust speed and gust direction changed those odds by at most 2 per cent, and two of the nine air terms differed from 1, wind in the eagle (0.98) and gust speed in the stork (0.98). Fitted per bird, with the four axes treated the same way, the median slope on turn rate was positive and differed from zero in all three species, while no air axis did in any of them. Updraft was left out because it is computed from turn rate and from the circle’s own climb, so it carries nothing about the air. Admitting it moves the turn’s odds ratio by under 2 per cent.

**Figure S7:**
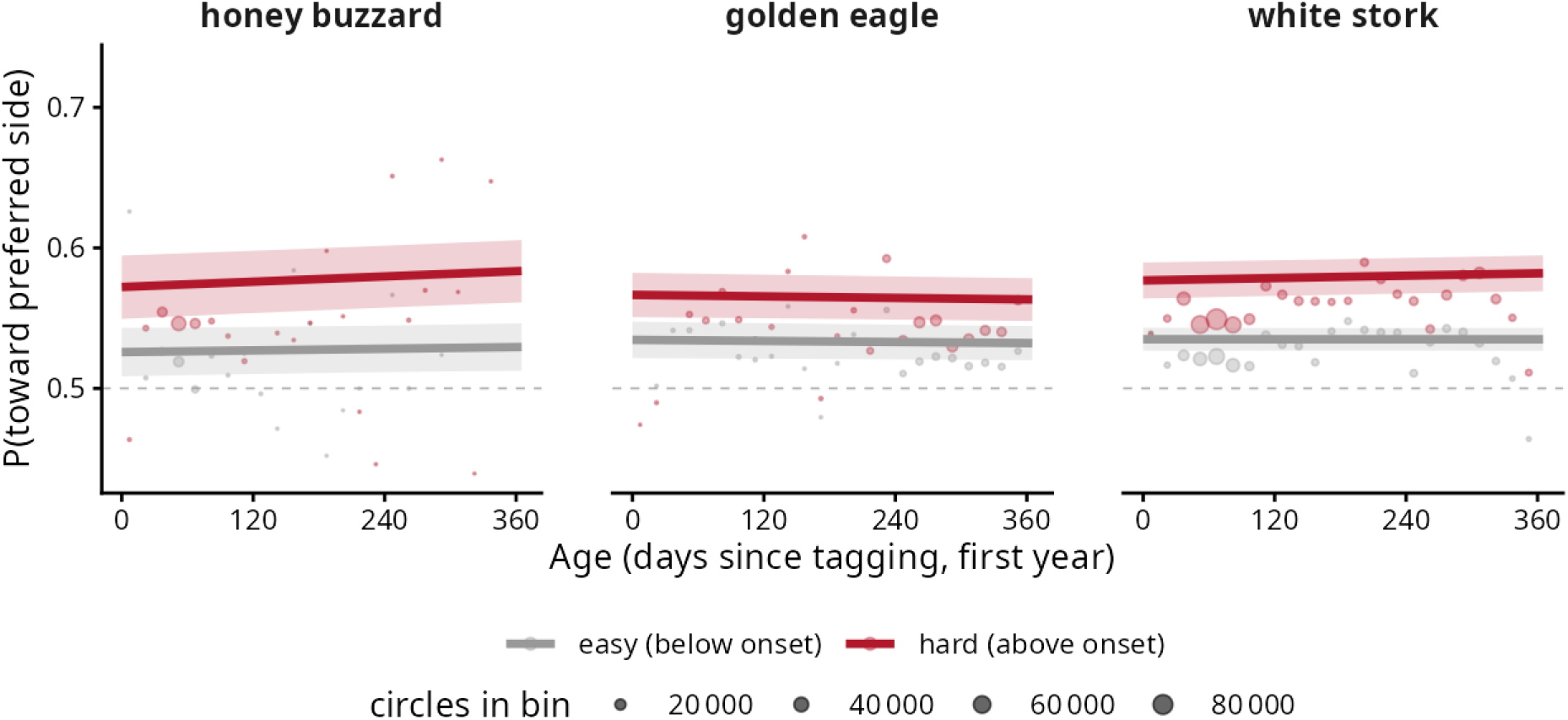
Laterality changes little with development. Fitted development-model predictions of the probability of turning towards each bird’s own preferred side, against age over the first year, for easy circles (below the onset, grey) and hard circles (above the onset, red). Bands are 95 per cent confidence intervals and points are the observed binned data behind the fit, sized by the number of circles in the bin. The curves are essentially flat: on hard circles the preferred-side probability rises by only about one percentage point over the whole first year in honey buzzards and about half a point in white storks, and does not change in golden eagles, with young and old confidence intervals overlapping throughout. The age-by-gain interaction is nonetheless statistically significant in the two migrants (p = 0.013 and < 0.001), significant only because it is estimated over hundreds of thousands of circles. Its magnitude is negligible, which is why development is reported as a minor result rather than a main-figure finding (main text, Results).

### S6: Life-cycle staging

Several questions condition on life-cycle stage: development, and affordability, by which we mean whether a bird expresses its side less when performance is at a premium, as in migration, and more when it is not. So the stage assignment must be independent of the turning data it is later crossed with, and robust to the imprecision of any automatic segmentation.

Stage is assigned by a latitude state-machine (summering, migration, wintering, resident) built from GPS alone, with no input from the turning data. Validated against curated stage dates for 31 honey buzzards (figure S8), the detected migration start is later than the curated departure in every bird, by a median of 17 days (the smoothing window trails a gradual departure, so stage analyses near the summering-to-migration boundary are biased conservatively). At the end of migration the detected date falls earlier than the curated one in most birds, the two differing by a median of 9 days.

**Figure S8:**
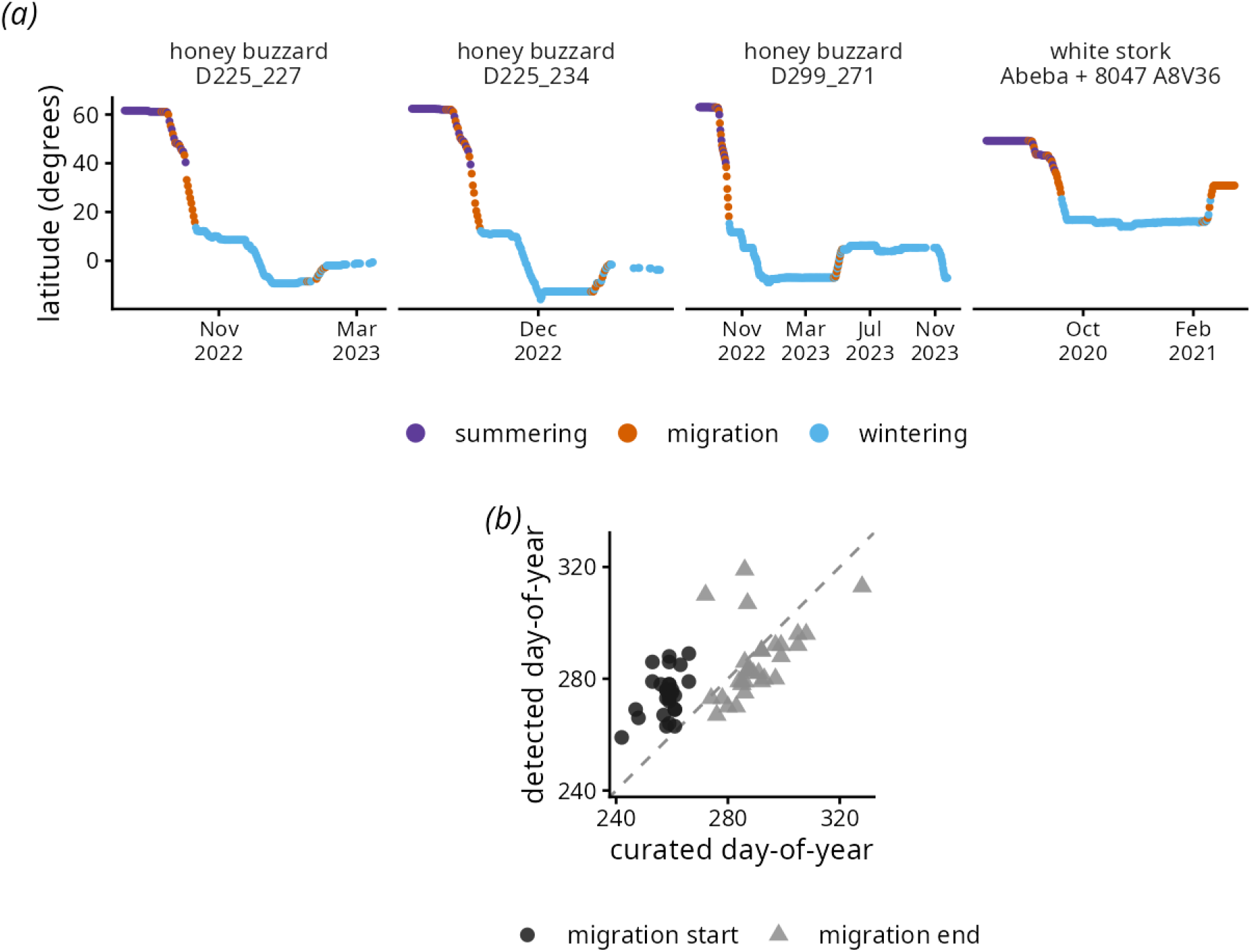
Life-cycle staging and its validation. (*a*) The latitude state-machine segments each bird’s annual cycle into summering, migration and wintering, from GPS alone and independently of the turning data. Four example migrants are shown, with daily latitude coloured by stage. (b) Validation against curated migration dates for 31 honey buzzards: detected migration start (circles) and end (triangles) day-of-year against the curated values, relative to the dashed line of equality. The detected start is later than the curated departure in every bird, by a median of 17 days (the smoothing window trails the gradual departure, so stage analyses near the summering-to-migration boundary are biased conservatively). At the end of migration the detected date falls earlier than the curated one in most birds, the two differing by a median of 9 days.

We quantified the effect of the lag: shifting every migration start by up to *±*17 days and every end by *±*9 days across a symmetric grid and refitting the stage-conditioned model at each node leaves the affordability verdict stable, with no migration trough at any node in either migrant. Here the bracketed ranges are the span of the log-odds stage contrast across the shift grid, not confidence intervals. In the buzzard both stage contrasts stay negative across the whole grid (post-fledging versus migration ranging from −0.11 to −0.01, wintering versus migration from −0.14 to −0.06), so migration remains the peak of expression. In the stork the wintering-versus-migration contrast stays positive and significant across the grid (0.07 to 0.17) while post-fledging-versus-migration hovers at zero. The lag moves the magnitude but never the qualitative conclusion.

For the buzzard specifically, the no-suppression reading rests on the well-determined wintering-versus-migration contrast, which survives Benjamini-Hochberg correction (log-odds contrast −0.133, p = 0.003, adjusted 0.005; table S3), not on the post-fledging-versus-migration contrast, which is only nominal and does not survive (log-odds contrast −0.089, p = 0.033, adjusted 0.053). Since the relabelling shifts every detected boundary, the reported range is an upper bound on the true single-boundary sensitivity, which makes the stable verdict only more convincing.

### S7: Residual independence and the population footnote

Circles within a thermal are serially dependent, which understates the standard errors of circle-grain estimates and makes their p-values too small. The headline is estimated between birds, so it does not carry that bias (figure S9).

**Figure S9:**
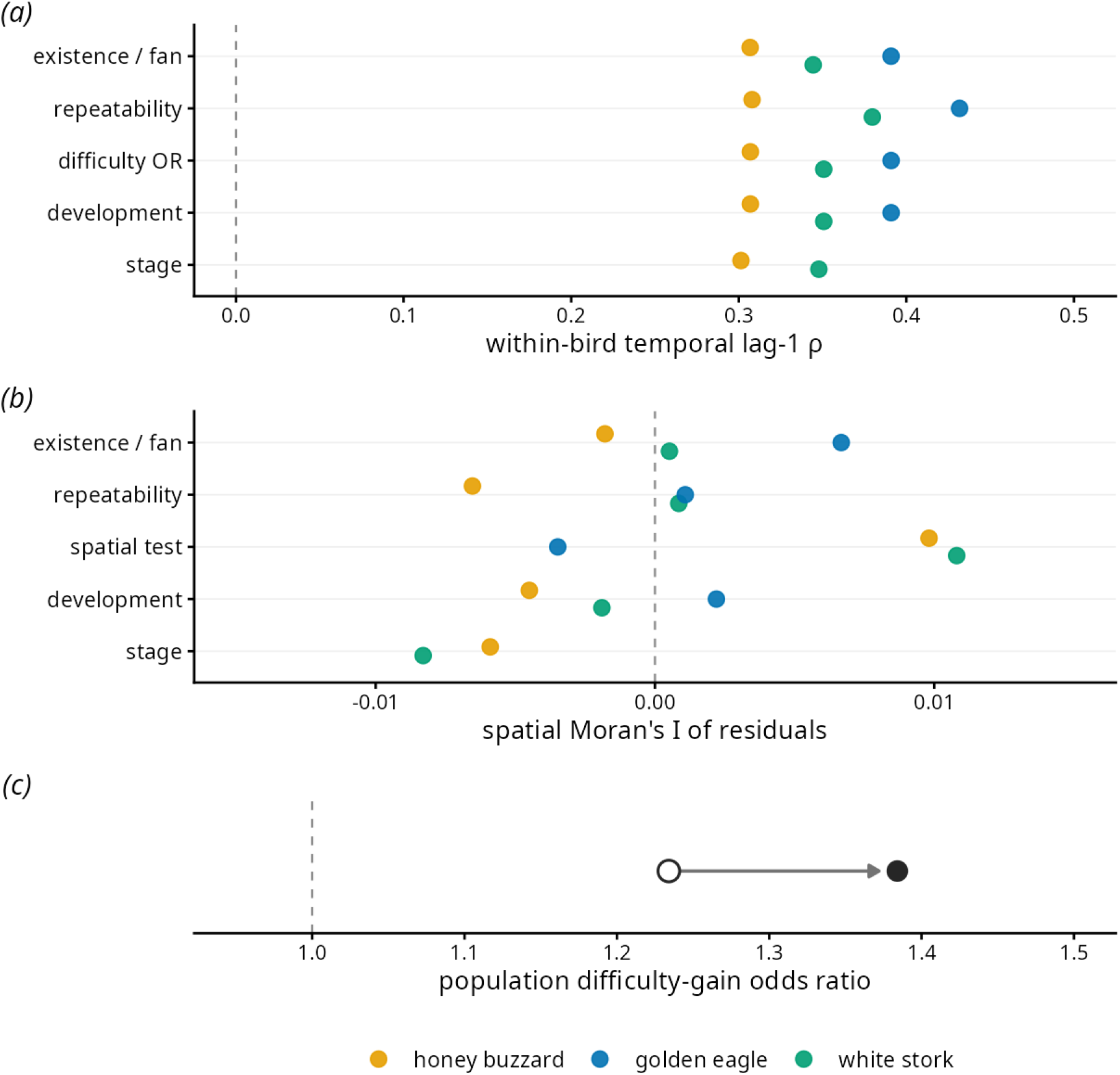
Residual independence and the population footnote. (*a*) The within-bird temporal lag-1 residual correlation is non-zero (about 0.3 to 0.4) in every inferential analysis, so it is checked and reported. Because the quantity estimated is at the level of the individual, it is not corrected. (*b*) The spatial Moran’s I of the residuals for the analyses shown is small (*|I| ≤* 0.011; the largest value anywhere, 0.14 in the IMU instability-level model, is outside the range drawn here) and, where formally significant, was shown not to move any estimate (see S7), so no spatial correction is applied. (*c*) An AR1-aware refit of the population difficulty-gain odds ratio, attempted per species as a sensitivity on the demoted population footnote, converged only for the honey buzzard, where it moves the keystone odds ratio from 1.23 (open circle) to 1.38 (filled), away from the null at 1 (dashed line), so the naive estimate is conservative. The golden-eagle refit did not converge and the white-stork refit converged only to a degenerate near-null. Neither is leaned on. The robustness of the estimate to autocorrelation rests on its between-bird random-slope structure, not on this footnote refit.

The constitutive dual residual check is run and reported per analysis (table S2). Spatial auto-correlation of the residuals (Moran’s I [S13]) is small in every analysis, *|I| ≤* 0.01 in the laterality models and at most 0.14 in the IMU instability-level model, but with 10^5^ to 10^6^ observations several of these small values are formally significant. Whether they matter was tested directly: adding an 11-km spatial grid random effect to the existence and difficulty models and to the climb-by-side model leaves each of those models’ own coefficients unchanged (difficulty slope on the raw side 0.142 vs 0.142 / 0.033 vs 0.033 / *−*0.103 vs *−*0.104; climb by side 0.0148 vs 0.0147 / 0.0010 vs 0.0008 / 0.0211 vs 0.0213, on this model’s sign convention rather than the bird-level contrast of table S2), so no spatial correction is applied. The live axis is temporal, where the within-bird lag-1 correlation is 0.31 to 0.39 in the laterality models and up to 0.55 in the IMU models. A correction is applied only where the check itself fails.

The headline is immune to this dependence because it is a between-bird quantity. The main model carries a random gain slope across birds ((1 + g_z | id)), so inference is over N independent birds, not over autocorrelated circles, and within-bird serial correlation cannot touch it. The only statistic that is susceptible is the population fixed-effect p, which is for that reason reported only as a check of parity. The check itself: in the main model (196 birds) the population probability of a rightward circle is 0.514 (p = 0.14) / 0.530 (p = 1 *×* 10*^−^*^6^) / 0.496 (p = 0.56). In the gain model on every bird with valid circles (248 birds) it is 0.515 (p = 0.048) / 0.518 (p = 0.002) / 0.490 (p = 0.069). The main text reports the main model. The population mean is flat below the onset by construction (the floor). Above it, the fixed slope of P(right) on difficulty is 0.14 [0.01, 0.27] (p = 0.038) / *−*0.01 [*−*0.07, 0.06] (p = 0.84) / 0.00 [*−*0.11, 0.11] (p = 0.98) per SD, so the honey buzzard’s population mean drifts rightward on tight circles (to 0.58 two SD above the onset), a composition effect of more right-than left-leaning birds in a cohort of 37, while the eagle holds its offset and the stork holds parity.

As a sensitivity on that footnote alone, an AR1-aware refit was attempted per species. It converged usefully only for the honey buzzard, where the keystone gain odds ratio moves from 1.23 to 1.38, away from the null, so the naive estimate is if anything conservative. The eagle refit did not converge and the stork refit reached only a degenerate near-null (*ρ ≈* 0, *p* = 0.89). Neither is leaned on, and the footnote’s robustness in those species rests on the between-bird random-slope structure, not on an AR1 correction.

### S8: Costs, in full

The claim that the bias is nearly free rests on null and bounded effects, so each is given with the interval that bounds it. The cost picture is given at three grains: the per-manoeuvre signature, the daily and seasonal integration, and a deterministic accounting that states the magnitude of “small” positively (figure S10).

**Figure S10:**
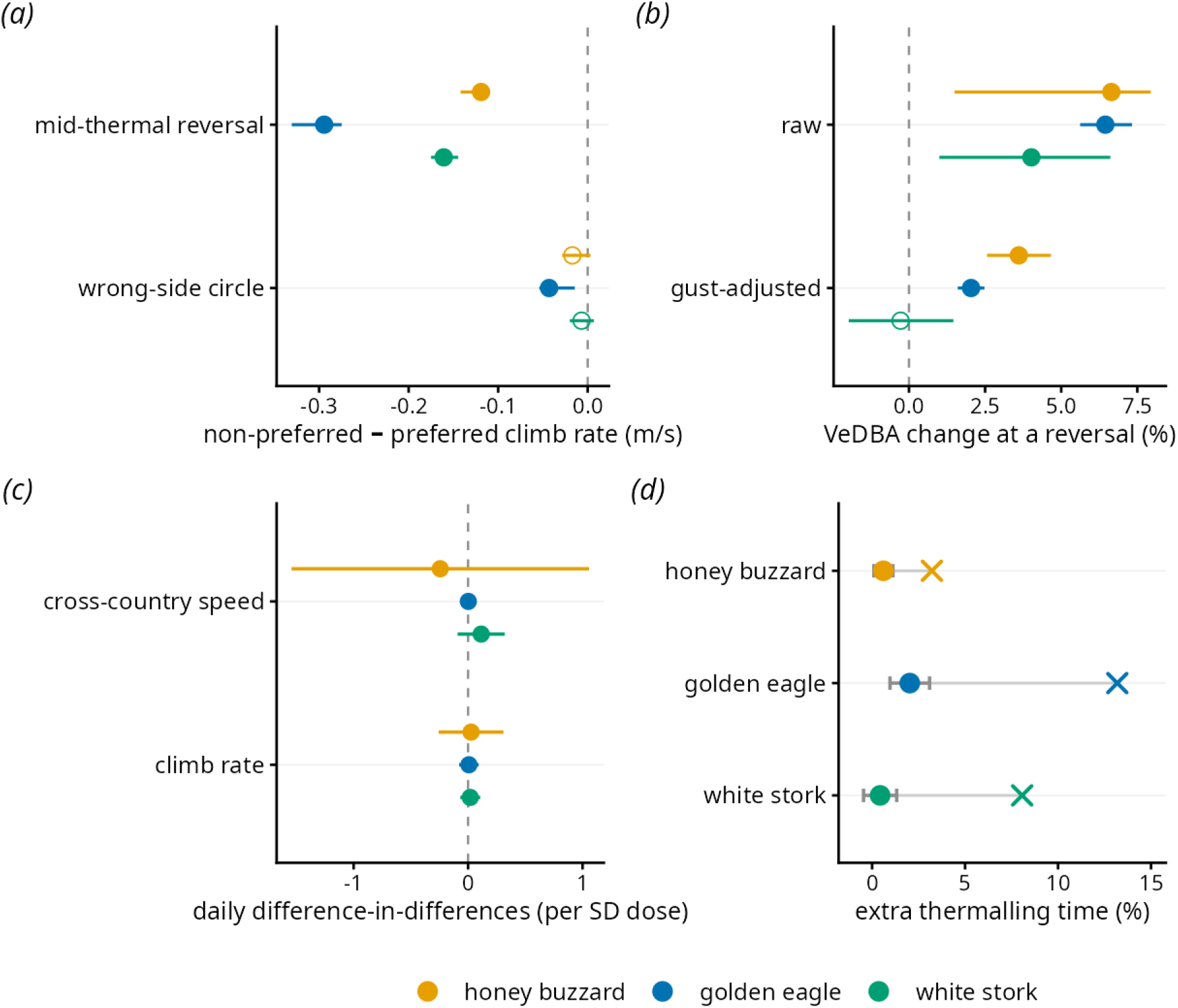
The costs. (*a*) The per-manoeuvre climb cost (bird-level within-thermal contrast, 95 per cent CI). In panels (*a*) and (b) a filled point is significant on the per-bird test and an open point is not. The mid-thermal reversal (switching) climb deficit is present and significant in all three species (filled, CIs below zero). The wrong-side (preferred versus non-preferred) climb deficit is significant only in the golden eagle (filled), with the honey buzzard and white stork non-significant (open, CIs spanning zero). (*b*) The real reversal effort spike (percentage increase in dynamic body acceleration, 95 per cent CI), raw and conditioned on gust. It survives gust conditioning in honey buzzards and golden eagles (filled) but not in white storks (open). (*c*) Yet the cost does not accumulate: the daily difference-in-differences of climb rate and cross-country speed (lateralised versus a symmetric placebo, within-bird, cluster-robust) spans zero in every species. (*d*) Accounting the estimated per-circle deficits onto realised behaviour bounds the extra thermalling at 0.6, 2.0 and 0.4 per cent of daily climbing time (filled, side-only, 95 per cent CI). The switch-inclusive upper bound (cross) is 3.2, 13.2 and 8.1 per cent.

At the per-manoeuvre grain the costs are real in the two raptors. On the per-bird test the non-preferred side is flown with more in-flight instability, turn-rate-matched; the mid-thermal reversal carries a climb deficit in all three species and an effort spike in the raptors; and the wrong-side climb deficit is significant in the eagle. The instability contrast is a within-bird paired comparison, so it includes only birds with IMU-recorded circles on both sides: all 37 buzzards and 27 eagles, but 54 of 58 IMU storks, since storks carry the fewest acts per bird and four contributed acts on one side only.

#### Is it better to be symmetric?

The per-bird instability level (the random intercept of a model of log instability on turn rate, gust and updraft, IMU subset) was compared between symmetric and lateralised birds. Medians differed by −0.003 / −0.033 / −0.051 on the log scale (symmetric minus lateralised, so a negative value means symmetric birds were marginally steadier), Wilcoxon p = 0.73 / 1 / 0.33, on 16 vs 19, 5 vs 19 and 7 vs 22 birds. The test is declared in the primary family and corrected within it (see S10). Its power is slight: the eagle and stork rest on five and seven symmetric birds, and the levels are estimated random effects entered without their uncertainty.

The kinematic circling-control signals cohere into one reading. The load factor (the ratio of lift to weight, which rises with bank angle) is slightly lower on the non-preferred side (side difference −0.0057 / −0.0030 / −0.0041, all p < 0.003): the bird banks more shallowly rather than over-banking. Read with the higher instability, this is a coherent motor-fluency signature: a tentative, under-committed bank must be nursed with continuous correction and circles wider, drifting the bird towards the weaker-lift thermal edge where it climbs marginally worse. We offer this chain as the natural interpretation of the joint pattern, not a demonstrated mechanism. It also bounds the climb-by-side result: because the same shallow geometry places the bird in weaker core updraft, the wrong-side climb deficit is confounded by radial position within the thermal, and we report it strictly as an association.

At the daily grain the cost does not integrate. The daily difference-in-differences (an extensive difficulty-weighted non-preferred-side count against a symmetric placebo, within-bird fixed effects, cluster-robust) is null on climb and cross-country speed in all three species, and its confidence intervals bound any daily side effect to a negligible range (climb DiD 0.03 [−0.26, 0.31] / 0.01 [−0.08, 0.09] / 0.02 [−0.07, 0.11] m s*^−^*^1^; cross-country −0.24 [−1.54, 1.06] / 0.00 [−0.05, 0.06] / 0.11 [−0.09, 0.32] m s*^−^*^1^). This is a null with a narrow interval, not absence of data. Eagles and storks flew less steadily on days when they circled more often against their preferred side (0.031, p = 0.004, and 0.030, p = 0.007, both surviving the correction in section S10). Their effort did not rise and their performance did not drop. The unsteadiness therefore shows up whether we count single manoeuvres or whole days, but it does not turn into a cost: of the five daily measures only the buzzard’s glide sink reached nominal significance (+0.18 [0.01, 0.35] m s*^−^*^1^, p = 0.038), and it does not survive the correction.

The buzzard daily DiD rests on about 37 clusters, the regime where the asymptotic cluster-robust interval is anti-conservative. The eagle and stork have on the order of ten thousand bird-days each and are unaffected. We therefore treat the buzzard daily interval as a lower bound on precision and do not rest the “does not accumulate” conclusion on the buzzard daily p. That conclusion is carried instead by the deterministic accounting below, by the well-identified eagle and stork daily DiDs, and by the per-manoeuvre costs that are significant in the raptors.

The deterministic accounting states the magnitude positively: integrating the estimated per-circle deficits onto each bird’s realised circle durations gives about 0.6 / 2.0 / 0.4% of daily climbing time from the side bias alone, a few minutes over a migration. A switch-inclusive figure (3.2 / 13.2 / 8.1%) additionally charges the reversal deficits, but those are reverse-causation-bounded and so a loose over-count, not a true upper bound. Each estimate is a no-compensation bound. Its interval is a propagation band, carrying the per-circle deficit’s bootstrap SE through an integration that holds each bird’s durations fixed, not a full sampling CI.

Two caveats complete the picture. VeDBA [S14, S15] captures the dynamic component of effort (instability, flapping and manoeuvring are genuine costs it registers), and the steady side difference in it is near-null (−0.9 / +0.4 / −0.7%, all ns), with the only positive signal a small transient spike at reversals, but VeDBA is insensitive to the isometric cost of holding a bank, so the ecological conclusion rests on the GPS climb, speed and migration results, which carry no such limitation. Migration performance versus laterality is reported as no detectable relationship, from a small-sample arm in the secondary family (n = 15 buzzards and 90 storks; eagles not testable) with a shrunken predictor biased towards the null, so it is not evidence of no relationship.

#### Wind, gust and updraft

Per circle, the horizontal wind (*u*, *v*) and the airspeed were estimated from the bird’s own circling with the thermal-soaring method of Weinzierl et al. [S4], airspeed being the magnitude of ground velocity minus wind. Circles on which the estimator diverged (wind above 40 m s*^−^*^1^) carry no wind. Gustiness is defined per thermal as the standard deviation of the estimated wind speed and the circular standard deviation of its direction across the thermal’s circles. Updraft is the vertical air speed, the observed climb rate plus the bird’s circling sink rate from the Pennycuick glide polar at the kinematic bank, and is used as a conditioning covariate only. For the IMU acts, which are not simultaneous with the GPS circles, the per-circle field was interpolated to each act’s position and time by per-bird space-time kriging, with the leave-one-out correlation of the wind interpolation computed per bird as a diagnostic of the interpolation.

### S9: Confounds and claims

A behavioural side-bias invites several alternative explanations. The alternatives considered are neuromuscular or sensory maturation, environmental context, movement mode, thermal availability and turbulence, route and geography, individual quality, sampling effort, and days-since-tagging as a coarse experience proxy. This section also states explicitly what is not claimed: neural lateralisation, innateness and causal learning are beyond what telemetry can establish and are not asserted.

#### Reverse causation

Turn rate is partly the bird’s own choice, and a bird flies its preferred-side circles slightly tighter than its non-preferred ones. The rise of the preference with turn rate therefore has two possible sources, the thermal’s demand and the bird’s choice, and a track cannot separate them: a bird holds one side for most of a thermal, so a thermal’s mean tightness carries the same choice. We state the gradient as descriptive. The causal reading rests on the IMU instability asymmetry at matched turn rate (see S8), which does not depend on which of the two set the turn rate. The trigger test (wind speed, gust speed and gust direction entered beside tightness; see S5) shows the preference following turn rate, with the air terms moving the odds by at most 2 per cent, whichever of the two set the turn rate.

### S10: Multiplicity

The study reports many tests across three species. We declared two disjoint families in advance and Benjamini-Hochberg-correct [S16] within each, reporting survivors and non-survivors alike.

The primary family is the headline inferential p per question and species across the difficulty, repeatability, development, switching and affordability tests, corrected over the whole family. Of 22 tests, 13 survive at FDR 0.05 (table S3): every difficulty-conditioning gain OR, all three within-thermal reluctance-to-leave tests, all three repeatability tests, the wintering-versus-migration affordability contrast, and the buzzard and stork age-development effects. The non-survivors are the experience-to-propensity null, the “is it better to be symmetric” null, the eagle age-development, and the post-fledging-versus-migration contrasts.

The within-thermal reluctance-to-leave effect enters the family at the bird level, not as a model Wald p over hundreds of thousands of transitions, which would be the circle-grain statistic S7 demotes. Per bird, the difference in switch rate between currently-on-non-preferred and currently-on-preferred circles is tested by a paired Wilcoxon across birds. It holds in all three species, the median bird-level difference being +0.025 / +0.023 / +0.030 over 37 / 71 / 124 birds. This is a reluctance to leave the preferred side, not a restoring force: of the switches that occur, the fraction returning to the preferred side is at chance (0.498 / 0.499 / 0.501).

The secondary “consequences” family (the per-manoeuvre and daily performance cost and null tests; the deterministic cost accounting has no p-value and is not part of the family) is corrected within itself (table S3b). Of 38 tests, 9 survive: the per-manoeuvre costs hold up (the switching climb deficit in all three species, the switching effort spike in the eagle, the wrong-side climb deficit and the instability asymmetry in the eagle, the instability asymmetry in the buzzard, and the daily instability-mechanism contrast in eagle and stork). The entire daily difference-in-differences tests of daily performance fail, and in particular the one nominally significant daily secondary, the buzzard glide-sink (raw p = 0.038), does not survive (adjusted 0.12). This settles the concern that a significant secondary was left uncorrected: within its declared family it is not significant.

**Table S3.**
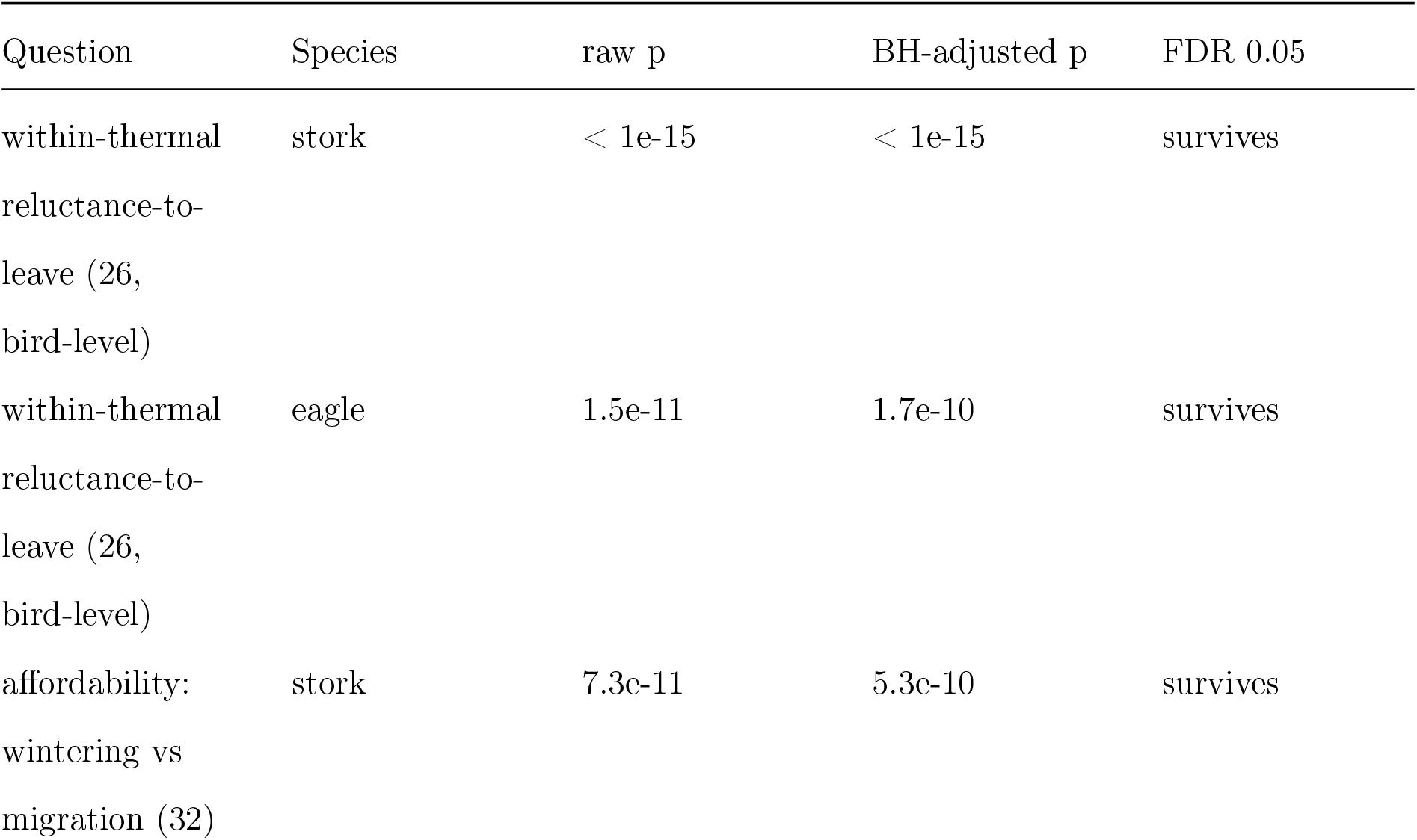
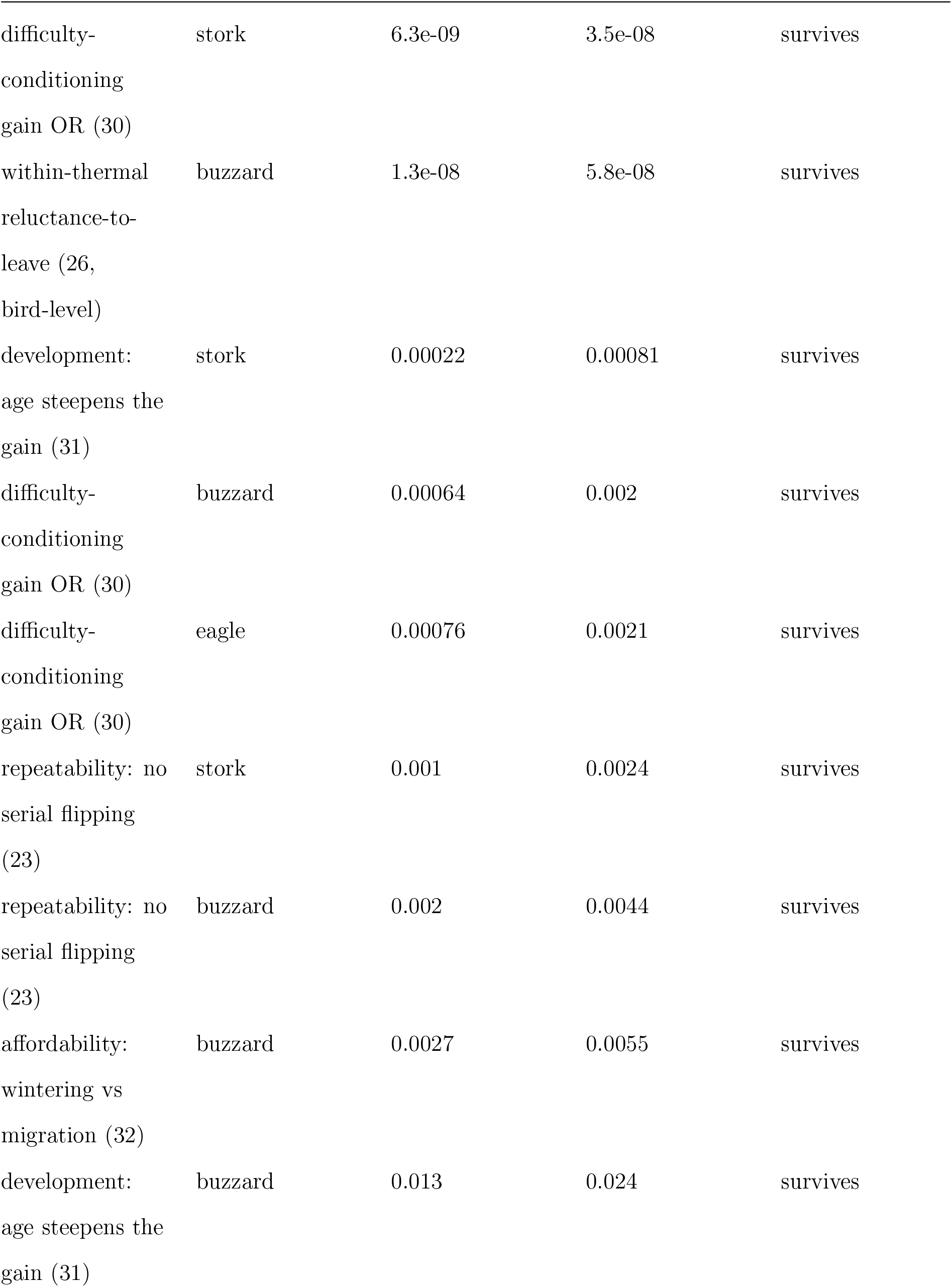
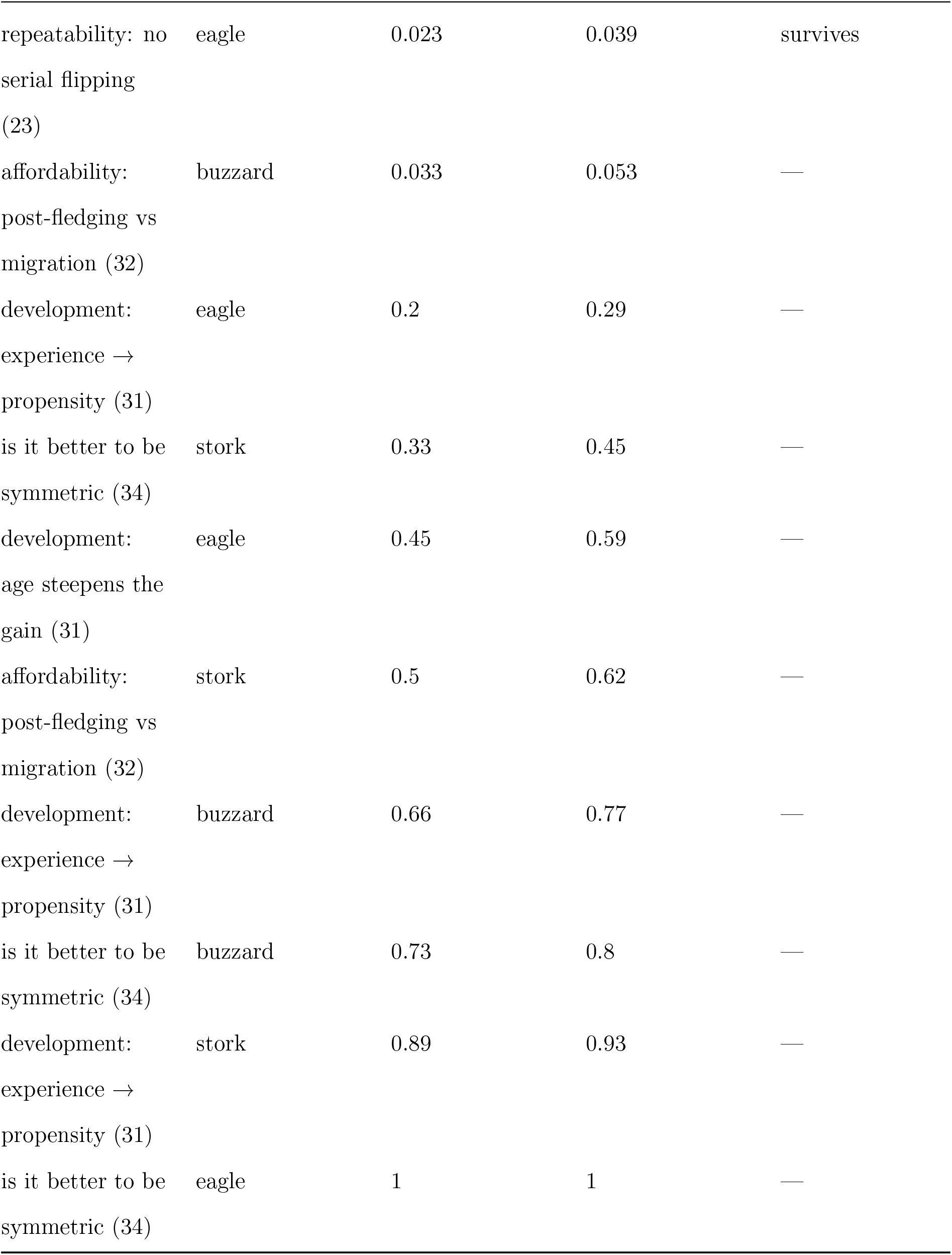
Benjamini-Hochberg multiplicity ledger. The 22 pre-specified primary-family tests (one headline inferential p per question and species), Benjamini-Hochberg-adjusted over the whole family; 13 survive at FDR 0.05, and the non-survivors are reported, not hidden.

**Table S3b.**
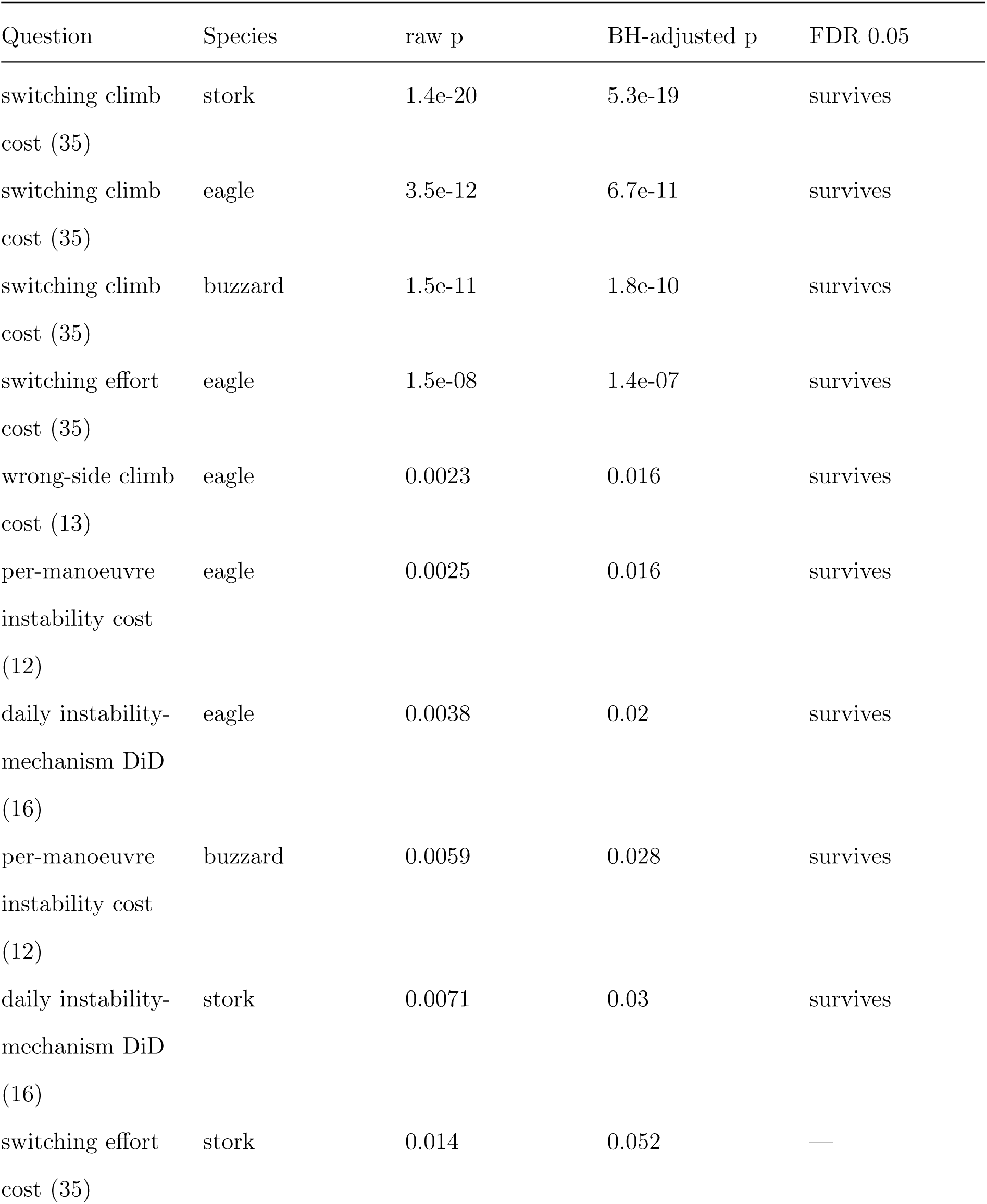
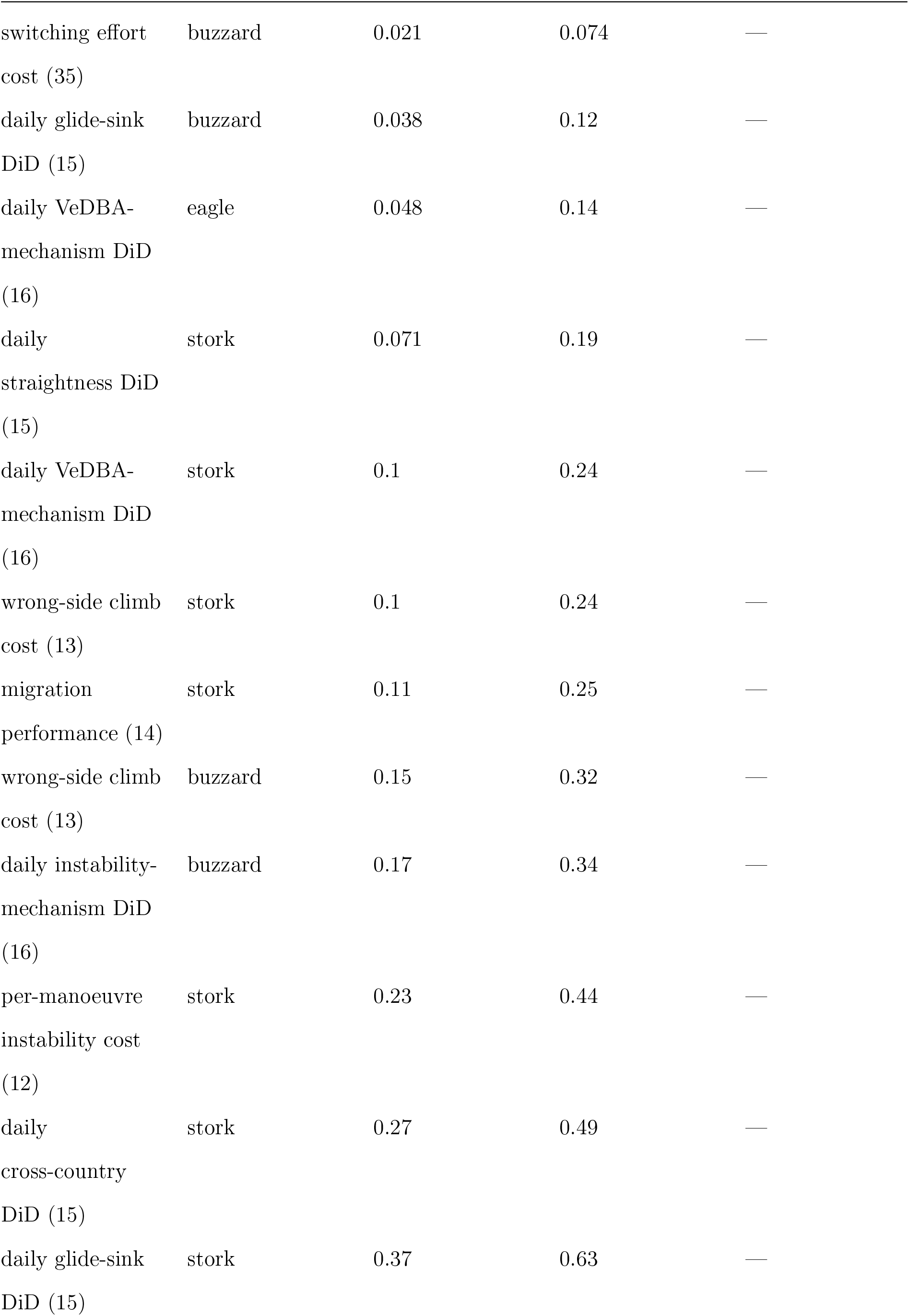
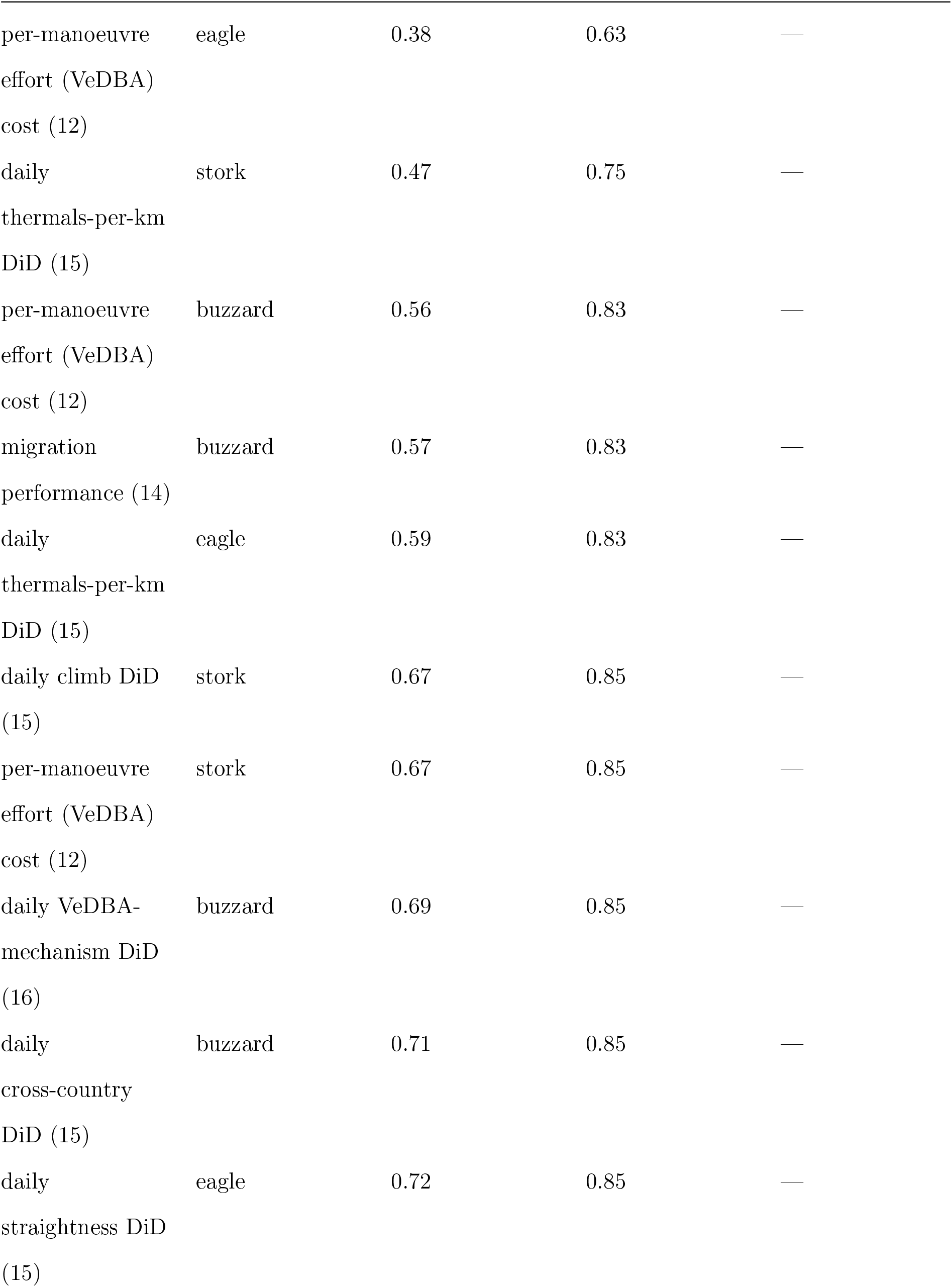
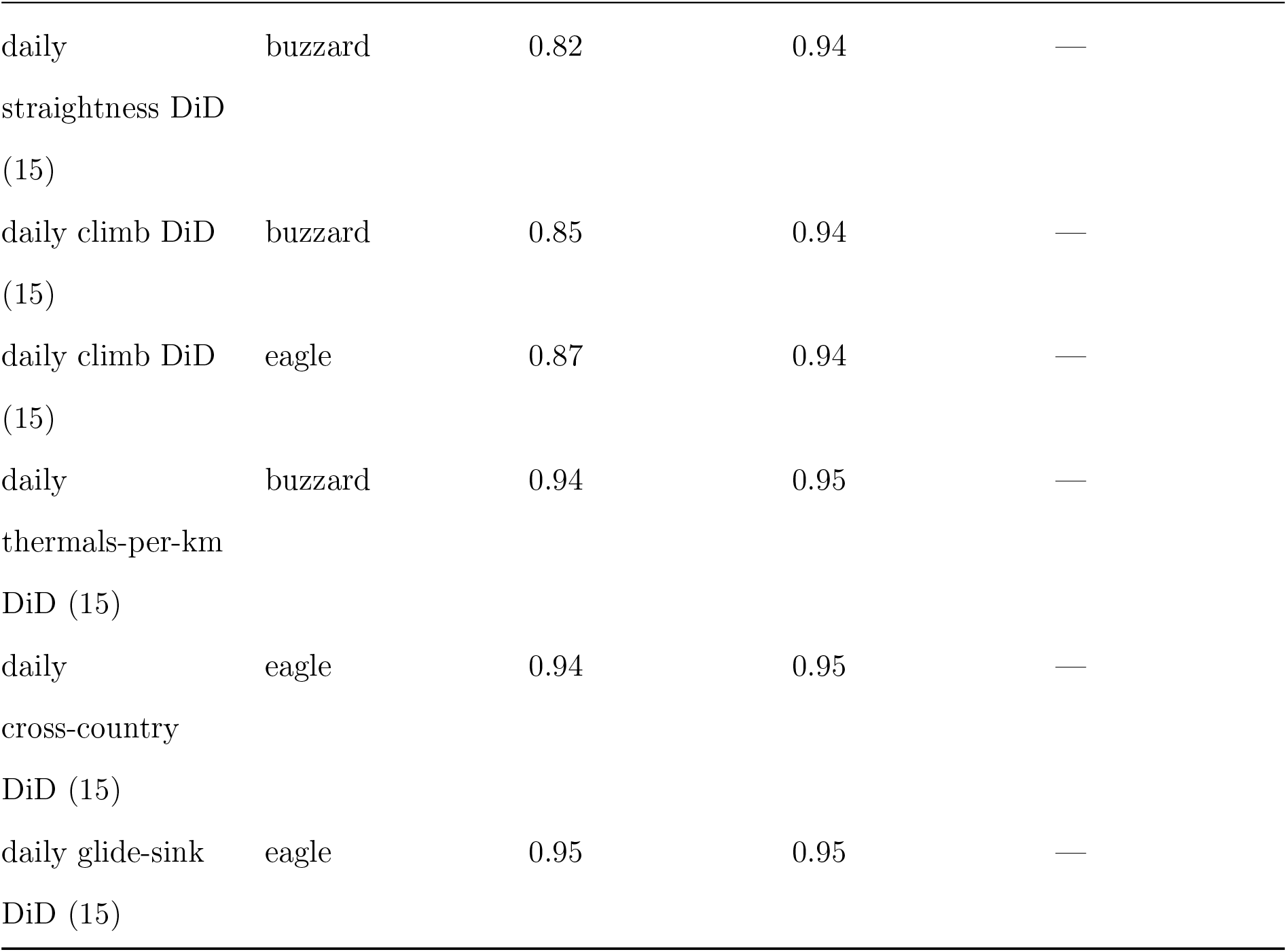
Secondary (consequences) multiplicity ledger. The 38 consequence tests (per-manoeuvre instability and effort, wrong-side and switching climb, switching effort, the daily difference-in-differences tests, the daily VeDBA mechanism and migration performance), Benjamini-Hochberg-adjusted within this declared secondary family; 9 survive at FDR 0.05. Reported for completeness: the claim that the bias does not accumulate rests on the bounded magnitude (the deterministic cost accounting) and the narrow confidence intervals of the daily nulls, not on uncorrected p-values, and the per-manoeuvre costs we report survive correction.

### S11: What a costly circle looks like, and how often birds circled together

Section S8 reads that the weak-side circle is flown wider and less steadily and therefore climbs worse. Its two end points are measured. The two links between them, that circling wider costs climb and that circling less steadily costs climb, are priced here. This section reports both, and then describes how often our tracked birds circled close to one another.

#### Why the obvious analysis cannot be run

Climb rate is the height gained divided by the circle’s duration, and both natural measures of how tightly a bird circles are ratios over that same duration: the turning rate is the net rotation divided by it, and the loop span scales with airspeed over rotation rate and tracks duration at a correlation of 0.85 within a thermal in every species. Regressing climb rate on either is therefore partly arithmetic. We measured how much: permuting the height gained among the circles of a thermal while keeping each circle’s own duration, which leaves only the shared denominator, still produces a slope 3.7, 1.9 and 2.4 times the one observed. The rate form has no power to answer the question and we did not use it. Instead we modelled the height gained per circle with duration, turning rate, loop span, side and position in the thermal on the other side of the equation, per bird and within thermals, so that nothing is a ratio and the thermal the bird was in is held exactly, and converted each bird’s coefficients to the climb currency afterwards using that bird’s own median circle duration.

#### Circling wider costs climb

At matched time in the thermal, each extra metre of loop span cost 0.039 [0.044, 0.032], 0.014 [0.016, 0.013] and 0.028 [0.029, 0.026] m s*^−^*^1^ of climb, and the coefficient was negative in 37 of 37, 70 of 71 and 109 of 110 birds. It survived a quadratic control for duration (0.037, 0.015, 0.027) and a response corrected for the height-for-speed exchange, which an altitude-only response would otherwise score as a climb loss (0.038, 0.017, 0.030). What this does not do is say why width costs climb. A wider loop at a given turning rate is a faster loop, so it carries more circling sink, and a bird drifting off the thermal core both widens its loop and meets weaker lift for a common reason. We claim that circling wide cost climb, not which of those delivered the cost, and we make no statement about where in the thermal a bird sat, which a track cannot establish.

#### Circling less steadily costs climb, where we can see it

On the circles that carry an overlapping inertial record, each natural-log unit of within-second in-flight instability cost 0.270 [0.422, 0.165] and 0.669 [0.932, 0.465] m s*^−^*^1^ of climb in honey buzzards and golden eagles, negative in 13 of 14 and 22 of 24 birds, and it survived the quadratic control for turning rate that section S2 establishes is necessary for raw in-flight instability (0.252 and 0.696). We could not estimate it in white storks. Only 2.2 per cent of stork circles carry an overlapping inertial act, against 10.9 and 17.5 per cent in the two raptors, and those few fall in short thermal fragments, so three birds had enough matched circles. That is a limit of sensor overlap and it is not evidence that the stork paid no such cost.

#### What is left of the side deficit

Holding the circle’s duration, turning rate, width and position in the thermal, the climb difference between the two sides was 0.014 [-0.027, 0.043], -0.010 [-0.043, 0.030] and 0.016 [0.004, 0.039] m s*^−^*^1^, so the deficit reported in section S8 did not survive holding the geometry of the circle in any species, and in the white stork the residual changed sign. This is a decomposition and not a direct effect. Width, turning rate and duration are themselves side-dependent on the account being tested, so holding them conditions on the very steps the account proposes, and a residual of either sign is what such conditioning can produce.

#### How often birds circled together

Among the moments when two tracked birds of the same study were both recording and both circling, they were within 100 m of each other in 0.002 [0.000, 0.012], 0.12 [0.04, 0.22] and 4.2 [3.2, 5.5] per cent of cases in honey buzzards, golden eagles and white storks, and within 1 km in 0.007 [0.000, 0.028], 0.20 [0.09, 0.32] and 5.2 [4.0, 6.7] per cent. Weighting every pair of birds equally rather than by how often it was seen gives 0.002, 0.17 and 2.7 per cent at 100 m. The stork figure rests on 575 of its 3 175 pairs of birds, the eagle’s on 34 of 1 127, and the buzzard’s on a single pair, so the buzzard value is an absence of observations rather than a measured frequency. Intervals come from resampling birds, since two birds flying together contribute many dependent observations. Three limits travel with these numbers. Only tagged birds are visible, so every figure is a lower bound and a bird recorded alone may have been among untagged companions. Proximity is a proxy for a shared thermal and not a detection of one, and with a median of 12 to 13 seconds between the two birds’ circles within a window, the smallest distances are at the resolution floor of the measurement. And the exposure is uneven in a way that differs between species: the densest tenth of days carries 93, 45 and 69 per cent of all pair observations, so the comparison is a description of our samples and not an estimate of a difference in behaviour.

#### What we do not claim

We do not report a comparison of these frequencies against chance. Five references were constructed and each was rejected on inspection, because a rate of co-occurrence carries the tag duty cycle, because a criterion of any bird being nearby rises with how many birds were recording at once, because displacing tracks across a year removes the annual cycle and so barely moves a resident while moving a migrant to another continent, and because a reference built by reading one bird against another some days later weights each pair of birds differently on its two sides. No test of whether sharing a thermal changes what a bird’s own circling is worth to its climb is reported, because thermals scored as flown alone necessarily include periods when no second tag was recording, so tag density would have been the treatment rather than a control. The frequency above is therefore offered as a property of our samples, and nothing in this section is a test or an explanation of why the per-manoeuvre costs resolve less clearly in the white stork.

